# Pharmacological Inhibition of BTK reduces neuroinflammation and stress induced anxiety *in vivo*

**DOI:** 10.1101/2021.01.11.426241

**Authors:** Simantini Ghosh, Zaidan Mohammed, Itender Singh

## Abstract

Stress related disorders lead to serious psychiatric disabilities and are comorbid with anxiety and depression. Current therapies targeting several neurotransmitter systems are only able to mitigate symptoms partially. It is well recognized that stress and trauma related disorders lead to a prominent inflammatory response in humans, and in several animal models a robust neuroinflammatory response has been observed. However, the therapeutic potential of targeting specific components of the inflammatory response has not been adequately studied in this context. The current study investigated the NLRP3 - Caspase1-IL-1β pathway, which recent research has identified as a major contributor to exacerbated inflammatory response in several peripheral and central nervous system pathological conditions. Using two different models of stress, first - single prolonged restraint stress followed by brief underwater submersion and second - predator odor exposure in mice, we demonstrate heightened anxious behavior in mice one-week after stress. Females in both models display an exacerbated anxiety response than males within the stressed group. Consistent with this data stressed animals demonstrate upregulation of IL-1β, IL-6, Caspase1 activity and NLRP3 inflammasome activation in brain, with female animals showing a stronger neuroinflammatory phenotype. Pharmacological inhibition of NLRP3 inflammasome activation led to a rescue in terms of anxious behavior as well as attenuated neuroinflammatory response, both of which were significantly more prominent in female animals. Further, we observed induction of activated Bruton’s Tyrosine Kinase (BTK), an upstream positive regulator of NLRP3 inflammasome activation, in hippocampus and amygdala of stressed mice. Next, we conducted proof-of-concept pharmacological BTK inhibitor studies with Ibrutinib, a drug that is already FDA approved for use in certain types of lymphomas and leukemias, as well as a second inhibitor of BTK, LFM-A13. In both sets of experiments, we found inhibition of BTK significantly reduced the anxious behavior in stressed mice and attenuated the induction of NLRP3 inflammasome, Caspase 1 and IL1β. Our results suggest that BTK inhibition can be further investigated in context of human stress and trauma related disorders as a therapeutic strategy.

## Introduction

Posttraumatic stress disorder (PTSD) is characterized by an impaired stress response comorbid with severe anxiety among other symptoms(Bryant, 2019). PTSD is a common finding after exposure to occupational and personal traumatic events leading to immense health and socioeconomic ramifications (Krystal et al., 2017). According to one estimate globally 345 million adult war survivors are likely to have experienced some PTSD symptoms (Hoppen and Morina, 2019), and this statistic does not take in to account any other trauma type, including sexual violence, which is often as potent a contributor to PTSD as war exposure (Friedman et al., 2011; Bryant, 2019). Present therapies of PTSD are often symptomatic, and fall short of treating PTSD and preventing relapse in long term (Taylor et al., 2003; Watts et al., 2013; Friedman and Bernardy, 2017; Krystal et al., 2017). With the advent of advanced research methods in neuroscience, the understanding of the neurobiology of PTSD is rapidly evolving. It is well documented that PTSD is associated with a chronic pro-inflammatory state, characterized by an elevated secretion of IL1β, IL-6, TNF-α, and other inflammatory mediators by the monocytes in the blood and microglia in the brain (Diering et al., 2016; Lindqvist et al., 2017; Mellon et al., 2018a; Sumner et al., 2020). The data on whether existing selective serotonin reuptake inhibitors (SSRI) or serotonin-norepinephrine reuptake inhibitors(SNRI) based pharmacological treatment regimens can ameliorate this neuroinflammatory phenotype is inconclusive at best (Tucker et al., 2004; Bonne et al., 2011; Sagarwala and Nasrallah, 2019). However, the therapeutic potential of targeting the chronically active neuroinflammatory pathways has not yet been investigated thoroughly. In this context, we set out to investigate the involvement of the multi-protein Nucleotide-binding Oligomerization Domain (NOD) Like Receptor (NLR) family, pyrin domain containing 3 (NLRP3) inflammasome complex in mouse models of stress. The NLRP3 inflammasome complex has been well described in the literature as pattern recognition receptors of the innate immune system expressed widely in the central nervous system and cells of myeloid lineage (Martinon et al., 2002; Agostini et al., 2004; Baroja-Mazo et al., 2014; Guo et al., 2015; Schroder and Tschopp, 2010). Inflammasomes evolved to act as host sensors to pathogen as well as endogenous stress, and work by the way of activating Caspase-1, ultimately leading to the production of the pleiotropic cytokine IL1β and IL18, leading to further neuroinflammatory sequalae that might compromise functionality (Halle et al., 2008; Jha et al., 2010; Geldhoff et al., 2013). IL1β in particular has been previously described as a major driver of the pro-inflammatory response in the brain and extensively investigated in the context of stroke as well as several neurodegenerative disorders (Heneka and O’Banion, 2007; Fann et al., 2013; Heneka et al., 2014; He et al., 2016). The contribution of NLRP3 activation towards pathophysiology and behavioral deficits in neurocognitive disorders and psychiatric disorder has emerged as an active area of investigation (Frank et al., 2016; Lénárt et al., 2016; Kaufmann et al., 2017; Song et al., 2017). In the present study, we demonstrate that there is a sexually divergent response in stress induced anxious behavior in mice subjected to either physical restraint stress with brief underwater submersion, or predator stress, alongside consistent upregulation of IL1β and IL6 expression in the hippocampus and amygdala at one-week post stress. We then investigated how pharmacological inhibition of NLRP3 activation alters neuroinflammatory markers in the hippocampus and amygdala of stressed versus controlled mice. Consistent with our hypothesis, and recent independent observation from other groups (Su et al., 2017; Dong et al., 2020a; Song et al., 2020), pharmacological inhibition of NLRP3 activation with a small molecule inhibitor MCC950 (Coll et al., 2015)reduced stress induced anxiety in mice and led to a reduction in hippocampal and amygdala IL1β as well as Caspase 1 activity. Additionally, we zeroed in on Bruton’s Tyrosine Kinase (BTK) as an upstream regulator of NLRP3 activation (Weber et al., 2017; Bittner et al., 2019). The overarching theoretical framework in which we operated was that once recruited, the NLRP3 complex initiates maturation of master regulator cytokines such as IL1β, which then directs a vicious cycle of self-synthesis and further neuroinflammatory reactions. Therefore, we worked with the hypothesis that if BTK could be inhibited, then we would see a rescue from stress induced anxiety with the reversal of the neuroinflammatory profile mirroring the data from the MCC950 experiment. To this end, we employed Ibrutinib, a BTK inhibitor, that is already FDA approved for human therapeutic use in certain types of leukemia and lymphomas (De Claro et al., 2015; Kapoor and Ansell, 2018). Ibrutinib treated mice showed marked reduction in stress induced anxiety and IL1β levels and Caspase 1 activity in the hippocampus and amygdala. We performed an orthogonal validation experiment with a second pharmacological inhibitor of BTK, LFM-A13, to observe more or less similar results.

## Material and Methods

### Experimental animals

Male and female wild-type C57BL/6 mice were obtained from Jackson Laboratories (Bar Harbor, ME). They were housed in vivarium under standard conditions (12 h: 12 h light-dark cycle starting at 7 am; 74°F; 55±10% humidity) in solid bottomed cages on woodchips bedding and had free access to autoclaved water and chow. All studies were performed according to the National Institute of Health guidance using protocols approved by Animal Studies Committees at Washington University School of Medicine and University of Delhi. Age-matched fourteen-week-old male and female C57BL/6 mice were used for experiments.

### Inhibitors

To pharmacologically inhibit NLRP3 inflammasome, a specific inhibitor MCC950 (Sigma-Aldrich, St. Louis, MO) was injected (IP: 50 mg/Kg) as reported in (Nair and Jacob, 2016), starting 72 hours before the induction of stress in mice, and again given at every 24 hours until the end of the experiment. Control mice were similarly injected with vehicle (0.05% DMSO in saline, i.p.).

To pharmacologically inhibit BTK, a specific inhibitor Ibrutinib (PCI-32765, Abcam, Cambridge, MA) was administered (IP: 3 mg/Kg) starting 72 hours before the induction of stress in mice, and again given at every 24 hours until the end of the experiment. Control mice were injected in parallel with the vehicle (0.05% DMSO in saline, i.p.) at the same time points. To validate the observation from Ibrutinib studies, a separate group of mice was injected (IP: 50 mg/Kg) with another BTK specific inhibitor LFM-A13 (Sigma-Aldrich) at similar time points as mentioned above. Control mice were similarly injected with vehicle (0.05% DMSO in saline, i.p.).

### Stress behavioral paradigm

Two weeks before behavioral testing, mice were gently handled daily. Mice were picked from the tail and permitted to explore freely on the gloved hand for 2 minutes. Mice were subjected to predator odor stress to induce psychological stress or restraint stress to induce physical stress. All stress induction experiments were performed at night between 8 pm and 3 am to avoid potential influence from noise and disturbances from day time activities.

To induce psychological stress, mice (pray) were placed away from the home cage in empty black plexiglass box (45 cm x 45 cm x 45 cm) containing fresh rat (predator) feces for 3 hours. The plexiglass box, as well as gloved hands, were wiped clean with 70% ethanol solution and let dry between tests with each mouse to remove any olfactory cues to mice. Since predator smell can come to the testing room and influence other animals; care was taken to expose the mice to rat feces under a fume hood.

To induce physical stress, mice were restrained by immobilizing them in disposable plastic tubes (4 cm – diameter x 12 cm – length) containing perforations near the nose for easy breathing and other ends with a hole in the closing cap for the tail to come out. Restrained mice were placed in a plexiglass box (45 cm x 45 cm x 45 cm) under the fume hood for 3 hours. Thereafter mice were immediately placed in a container with ice-cold water and allowed to swim for 45 seconds, and then gently submerged completely underwater for 30 seconds. Restraining plastic tubes were used only once for each mouse. The plexiglass box as well as gloved hands were carefully cleaned with 70% ethanol solution and let dry between each session.

Control mice were placed similarly in separate plexiglass boxes (45 cm x 45 cm x 45 cm) near the stress experiment arena, however, away from the fume hood. Control exposures were completed before starting the stress experiments to avoid any potential disturbance to the control of mice from the stress paradigm. As an additional control and to see the potential influence of control mice from the surrounding in the stress experiment arena, a set of control mice were kept in a vivarium away from the stress experimental arena.

### Behavioral tests

Seven days after the induction of predator odor and physical stress, the stressed and non-stressed control mice were examined for anxious behavior using open field test (OFT), light-dark test (LDT), and elevated plus maze test (EPM). All behavior assessment experiments were performed at night between 8 pm and 3 am when rodents are most active - being nocturnal, and also to avoid potential influence from noise and disturbances from day time activities. Between tests with each mouse, all the behavior tools and hands were thoroughly wiped clean with 70% ethanol and dried well to remove any olfactory cues from the experimental mice. Behavior tests were performed in the following sequence: open field test, light-dark test, and the elevated plus-maze with 1-hour interval between each test. These tests represent a standard battery utilized for the examination of anxious behavior (Bourin, 2015; Belovicova et al., 2017).

### Open Field Test

Examination of anxious behavior of stressed and non-stressed control mice was conducted using an open-aired rectangular gray-colored opaque box (30 cm x 45 cm) surrounded by a 35 cm high wall. At the beginning of the test, each mouse was placed into the same left spot at one corner of the rectangular arena and allowed to freely explore the arena for 7 minutes. The time spent in the central arena – marked by an 8 cm x 12 cm rectangle was measured to assess anxiogenic behavior. Stressed animals tend to show hesitation to spend time in a central area away from the sidewalls of the box (Prut and Belzung, 2003). The testing arena was cleaned with 70% ethanol before and between each trial.

### Light-Dark Test

This test utilizes a light-dark box to examine the innate tendency of mice to avoid the brightly lit area and escape to the safety of a dark area, especially when stressed (Crawley and Goodwin, 1980; Teegarden, 2012). The test box consisted of a rectangular plexiglass box (50 cm x 30 cm) with one-half enclosed with black opaque plexiglass sheet to serve as a dark area and the other half had a clear plexiglass enclosure to act as a light area. These two areas were connected with an opening of 8 cm x 8 cm to enable the mouse to freely move in these areas. Mice were let in the box for 10 minutes and time spent in each chamber was counted to assess the anxiety behavior in mice. The light-dark box was cleaned thoroughly with 70% ethanol between each trial to remove any olfactory cues for the mice.

### Elevated plus-maze

After OFT and LDT, mice were subjected to elevated plus-maze tests based on previously published methods (Carola et al., 2002; Carobrez & Bertoglio, 2005). Elevated plus-maze was made of four perpendicular plus-shape arms of plexiglass (30 cm in length, 7 cm in width), extending from the central square (7cm^2^) at the elevation of 60 cm from the floor. One set of opposing arms had 15 cm high walls – representing a secure area, and the other set of opposing arms were without walls – representing an unsafe area. Elevated plus-maze is based on the innate aversion of mice to open elevated unsafe spaces, and uses the conflict between exploration and this aversion. Mice were introduced into the central square of the elevated maze and allowed to freely explore all the arms for 5 minutes. Anxious behavior was determined by the percent time spent in the open arm of the elevated plus-maze. An arm entry was counted when all the four limbs of the mouse enter into the open arm. Increased time spent in the open arm denotes a lower degree of anxiety in the rodents (Carola et al., 2002).

### Tissue harvesting

Following the behavior tests, mice were anesthetized by injection with a mixture of ketamine (100 mg/kg) and xylazine (10 mg/Kg) and transcardially perfused with ice-cold heparinized phosphate-buffered saline (PBS) to clear the brain vasculature of blood and peripheral immune cells (Bell et al., 2012; Deane et al., 2012). The brains were quickly extracted, hemi-dissected, and hippocampus and amygdale isolated under the stereomicroscope. Tissues were snap-frozen in liquid nitrogen and stored in a −80°C freezer for later use.

### Quantification of IL1β and IL6

The snap-frozen mouse brain tissues were analyzed for IL1β and IL6 based on our published studies (Ghosh et al., 2013; Singh et al., 2013). Briefly, one-half of the hippocampus and amygdala were first homogenized with pellet pestle in RIPA buffer containing protease and phosphatase inhibitors (Thermo Fisher Scientific, Waltham, MA). These were further homogenized by sonication (Mesonix Sonicater 3000) using six pluses of 20 seconds each on ice. The homogenates were agitated on ice for 30 minutes and then centrifuged at 14,000 x g at 4°C for 30 minutes. The supernatant was immediately used to determine the levels of IL1β and IL6 using Quantikine Mouse ELISA kits and following the manufacturer’s protocols (R&D Systems, Minneapolis, Mn).

### Immunoblot analysis

Brain tissue homogenates in RIPA buffer containing protease and phosphatase inhibitors were mixed with NuPAGE LDS sample buffer (Thermo Fisher Scientific) and immediately proteins (10 µg) were separated by SDS PAGE electrophoresis (Bio-Rad Laboratories, Hercules, CA) on 4-12% Bis-Tris gradient gels (Thermo Fisher Scientific). The proteins were transferred onto the PVDF membrane (Bio-Rad Laboratories). The membranes were blocked in 5% w/v bovine serum albumin (IgG-free and protease-free BSA, Jackson ImmunoResearch Laboratories, West Grove, PA) in 50 mM Tris-HCl (pH 7.4), 150 mM NaCl and 0.1% Tween 20 (Tris-buffered saline with Tween, TBST). The membranes were incubated at 4°C overnight with the following mouse-specific primary antibodies: anti-NLRP3 (D4D8T, rabbit monoclonal antibody, 1:1000, #15101, Cell Signaling Technology, Danvers, MA), anti-cleaved Caspase 1 (p20, Asp296, E2G2I, rabbit monoclonal antibody,1: 1000, #89332, Cell Signaling Technology), anti-phospho-Btk (Tyr223, D9T6H, rabbit monoclonal antibody, 1:1000, #87141, Cell Signaling Technology), anti-Btk (D3H5, rabbit monoclonal antibody, 1:1000, #8547, Cell Signaling Technology) and anti-alpha tubulin for loading control (ab4074, rabbit polyclonal antibody, 1:5000, Abcam). The membranes were washed five times with TBST and incubated with horseradish peroxidase-conjugated goat anti-rabbit IgG secondary antibody (adsorbed with mouse and bovine to prevent cross-reactivity, 1:10,000, #STAR124P, Bio-Rad Laboratories) in TBST for 1 hour at room temperature. The membranes were washed five times with TBST and immunoreactivity was detected using SuperSignal West Pico PLUS Chemiluminescent substrate (Thermo Fisher Scientific).

### Caspase 1 activity assay

The activation of Caspase 1 in mouse brain samples were determined using Caspase 1 Colorimetric Assay Kit (YVAD, Merck Millipore, Burlington, MA) and following the manufacturer’s instructions. The assay determines the activity of Caspase 1 that recognizes the sequence YVAD. In brief, hippocampus and amygdale tissue samples from one brain hemisphere were homogenized in an ice-cold lysis buffer. The lysates were added to a Caspase 1 reaction buffer in a 96-well flat-bottom microplate. A substrate solution containing YVAD conjugated to chromophore p-nitroanillide (pNA) was added to each well and this was followed by incubation at 37°C for 2 hours. The quantification of Caspase 1 activity was carried out by the detection of pNA from the cleavage of the pNA-YVAD substrate by measuring light emission at 405 nm using a microplate reader (Molecular Devices, San Jose, CA).

### Statistical analysis and reporting

All analysis was carried out using the open source program JASP (JASP Team 2020, Version 0.14). All main effects are reported with eta square (η^2^) and partial eta square (η_p_^2^) for one-way and Factorial ANOVA tests respectively, to provide information about effect sizes for each test. In factorial ANOVAs, if interaction terms were significant, the main effects were not tested post-hoc. Instead interactions were broken down using proper Bonferroni corrected tests.Alongside p values, the t values are reported instead of Cohen’s d as measures of effect size in each case, since the Cohen’s d is not corrected for multiple comparisons in JASP. In case of 2X2X2 factorial ANOVAs, for each analysis, the results are reported with the significant main effects and the interaction term that had the largest η_p_^2^. If, however, the 3 way interaction term was statistically significant, then this was reported and post-hoc testing was done accordingly. P<0.05 was considered statistically significant for all analysis. Unless specified, all data reported in the text is reported as Mean ± SD.

## Results

### Both predator odor stress and physical stress leads to hyper anxious behavior in mice and elevated neuroinflammatory markers in the brain at 1-week post-stress

We conducted two different stress paradigms, one was psychological stress and another was physical stress, as described in the methods section. Psychological stress was emulated by exposing fourteen-week-old mice to fresh rat feces for three hours away from home cage. Rats are natural predators for mice and their odor is well known to induce anxiety behavior in mice (Hebb et al. 2003; Rorick-Kehn et al. 2005). To impart physical stress, we used a combination of physical restraint, forced swim and underwater submersion, as discussed in the methods section. We assessed anxious behavior 7 days post stress and soon after analyzed the animals’ brains for neuroinflammatory markers. To control for stress from the surroundings in the stress experiment arena, we used two different sets of control mice. The first group of control animals were brought into the stress experimental arena to expose them to the surroundings, however, kept away from the fume hood where stress experiments were carried out to avoid the confound of any vicariously acquired stress. The second group of control rodents was kept in the vivarium away from the stress experiment arena.

Animals exposed to predator odor and subjected to physical stress demonstrated significantly reduced crossings of the central field in an open field test, a behavior that is associated with increased anxiety in rodents (Figure 1A). A one-way ANOVA revealed a significant effect of the experimental stress manipulations [F (*3,78*) = 42.16, p<0.001, η^2^= 0.619; Figure 1A]. Pairwise comparisons with the Bonferroni correction demonstrated that the mice in the physical stress group performed worst (Mean ± SD: 13.9 ± 5.24), followed by the mice exposed to predator odor (21.7± 5.18), compared to both sets of control mice (p<0.001). The control mice from vivarium (32.67 ± 8.38)or in the stress arena with no direct view of the stress paradigms (33.57 ± 6.74) were not significantly different from one another in the time spent in the central square of the field (p >0.05).

**Figure 1.**
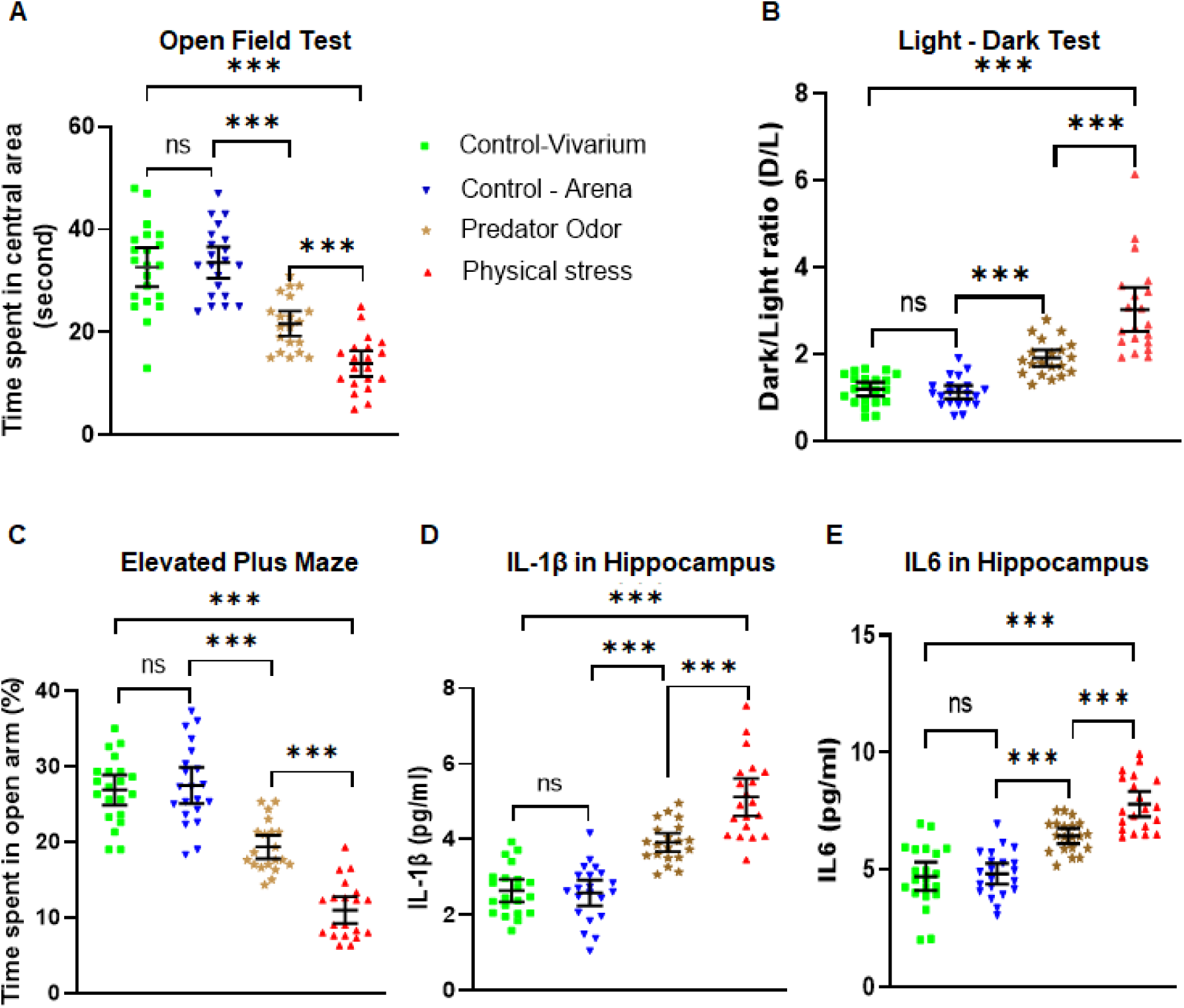
Physical and psychological stress induces hyper anxious behavior in mice. **(A)** Examination of anxious behavior of stressed and non-stressed control mice using open field test (OFT). Mice exposed to predator odor and physical stress exhibited significantly higher anxiety levels when compared to the controls (Control-vivarium, and Control-arena), as depicted by their hesitation to explore or spend more time in the central area of the OFT. **(B)** Light-Dark test (LDT) revealed mice exposed to predator odor and restrain stress exhibited significantly higher anxiety levels (Dark-Light ratios) when compared to the controls, as explained by their reluctance to spend more time in the light chamber of the LDT, with physically stressed mice displaying greater anxious behavior. **(C)** Elevated plus maze (EPM) test revealed mice subjected to predator odor, and physical stress displayed significantly increased anxiety levels compared to the controls, as evidenced by their hesitation to spend more time in the open arms of the EPM. **(D)** Examination of Interleukin 1β (IL1β) in the hippocampus of mice exposed to predator odor and physical stress showed aberrantly higher IL1β levels compared to the controls.**(E)** Examination of Interleukin 6 (IL6) in the hippocampus of mice by ELISA showed that mice exposed to predator odor and physical stress showed aberrantly higher IL6 levels relative to the controls. All data are presented as Mean with 95% CI (n=20-21/group); ***p<0.001, ns (not significant); one-way *ANOVA* (Shapiro-Wilk’s, *p*>0.05) followed by Bonferroni *post-hoc* test.

The second behavior test to access the anxiety in mice was the light-dark test. Naïve mice spend roughly equivalent amount of time exploring both the light and the dark compartments of the testing chambers with a slight preference on the dark compartment. A tendency of spending more time in the dark half of the testing chamber is associated with increased anxiety in mice. The ratio of the time spent by the mice in the dark to the time spent in the lighted half of the testing chamber (D/L ratio) should therefore be approximately 1 or slightly above 1 in control mice, and an increase in this ratio would reflect increased anxious behavior in mice. A one-way ANOVA of the experimental groups revealed a significant difference in group means by experimental status [F (*3,78*)=41.54,p<0.001,η^2^= 0.615; Figure 1B].

Bonferroni corrected post-hoc tests were used for pairwise comparisons. While both the control groups of mice, i.e. one in the vivarium and another exposed to the stress experiment arena displayed similar D/L ratios (1.19 ± 0.34 and 1.13 ± 0.36;*t*=0.37, p<0.05), mice exposed to predator odor exhibit a significant increase in the D/L ratio (1.92 ± 0.41) compared to control mice in the arena (*t*=4.06, p<0.001) and control mice in the vivarium (*t*=3.69, p<0.001). Mice subjected to physical stress exhibited the highest D/L ratio (3.03 ± 1.08), significantly more compared to mice exposed to predator odor, (*t*=5.72, p<0.001), control mice in the arena (*t*=9.85, p<0.001), and control mice in the vivarium (*t*=9.49, p<0.001).

To further evaluate the anxiety behavior in stressed mice, we conducted the well-established elevated plus maze (EPM) test. Time spent in the open arm of the maze was used to measure anxious behavior, with a reduction in the time spent in the open arm taken as a manifestation of increased anxiety. A one-way ANOVA revealed a significant difference among group means [F (*3,78*) =67.13, p<0.001,η^2^= 0.721; Figure 1C]. Post hoc testing with the Bonferroni correction was used for pairwise comparisons. Mice subjected to physical stress spent the shortest time in the open arm of the maze in seconds (33.05 ± 11.35), compared to both groups of control mice in the stress experiment arena (82.54 ± 15.01; *t*=12.38, p<0.001) and in the vivarium (80.71 ± 13.07; *t*=11.93, p<0.001), as well as the mice exposed to predator odor (58.1 ± 9.98; *t*=6.2, p<0.001). Mice exposed to predator odor also displayed significantly higher anxiety manifested by the shorter times they spent in the open arm of elevated maze, as compared to the control mice near the arena (*t*=6.114, p<0.001) and the vivarium (*t*=5.661, p<0.001). The two control groups were not significantly different from one another in this regard (*t*=0.5, p=1.0).

The above experiments demonstrated that the stress paradigms we conducted on mice elicited a robust anxiety response, and predictably, a higher response in mice subjected to physical stress. Next, we focused on the neuroinflammatory markers of the brain, specifically the pro-inflammatory cytokines IL1β and IL6 in the hippocampus. An ELISA assay for IL1β was conducted and analyzed by one-way ANOVA. Both physical stress and predator odor stress elicited a robust IL1β upregulation in the hippocampus of the mice, compared to control hippocampi [F (*3,78*) =49. 99, p<0.001,η^2^= 0.658; Figure 1D]. Post-hoc analysis (Bonferroni corrected) revealed that physical stress led to the most robust upregulation of IL1β (pg/ml) in the hippocampus (5.12 ± 1.06), compared to both groups of control mice in the stress experimental arena (2.58 ± 0.75; *t*=10.54, p<0.001) and in the vivarium (2.64 ± 0.65; *t*=10.29, p<0.001), as well as the mice exposed to predator odor (3.92± 0.53; *t*=4.92, p<0.001). IL1β levels in mice exposed to predator odor were significantly higher as compared to the control mice near the arena (*t*=5.59, p<0.001) and the vivarium (*t*=5.3, p<0.001). IL1β levels were similar between the two control groups (*t*=0.25, p=1.0).

IL6 levels in the hippocampus followed a very similar pattern to that of IL1β. A one-way ANOVA revealed significant differences in group means based on experimental conditions [F (*3,78*) =43. 79, p<0.001, η^2^= 0.599; Figure 1E]. Physical stress upregulated IL6 most robustly (7.79 pg/ml ± 1.15), compared to both groups of control mice in the arena (4.82 pg/ml ± 0.97; *t*=8.95, p<0.001) and in the vivarium (4.71 pg/ml ± 1.32; *t*=9.29, p<0.001), as well as the mice exposed to predator odor (6.43 pg/ml± 0.69; *t*=4.06, p<0.001). M ice exposed to predator odor displayed significantly elevated IL-6 compared to the control mice near the arena (*t*=4.84, p<0.001) and the vivarium (*t*=5.18, p<0.001). The two control groups were not significantly different from one another in their IL6 expression levels in the hippocampus (*t*=0.34, p=1.0).

Taken together, the results of the experiment suggested that alongside anxious behavior, our stress models also upregulated pro-inflammatory cytokines IL1β and IL6 in the brain. Physical stress elicited a more prominent response across all behavioral as well as molecular measures. Since the two control groups had no significant difference in any of the outcome measures, for the rest of this study, we conducted our experiments with control mice which were exposed to the stress experiment arena without visual access to the area where stress paradigms were performed.

### Female mice display an exacerbated stress response compared to male mice

To understand the gender differences in anxiogenic response, we examined the male and female mice under both stress paradigms, i.e. psychological stress (predator odor) and physical stress (restraint stress followed by underwater trauma). We observed that female mice had a more pronounced response to stress in both the stress models used, across behavioral and molecular outcome measures. However, like observed in the earlier experiments, physical stress continued to elicit a more robust response in all outcomes assessed compared to predator odor stress. The sex difference is of interest and special relevance because this mirrors human data on PTSD and stress and anxiety related disorders – females demonstrate a higher incidence and prevalence and severity of anxious symptoms in these disorders (Olff 2017).

Physically stressed mice, both male and female were assessed by the open field test (Figure 2A), the light-dark test (Figure 2B) and the elevated plus maze test (Figure 2C). We conducted separate analysis between groups using factorial ANOVAs for each test, with stress (Control mice near the arena vs. physically stressed mice) and sex (male vs. female) as factors. For the open field test, there was a significant main effect of stress [F (*1,64*) =294.76, p<0.001, η_p_^2^= 0.82]and a significant main effect of sex [F (*1,64*) =13.97, p<0.001, η_p_^2^= 0.17] on the time spent in the central square of the open field. This was further qualified by a significant interaction between the two factors [F (*1,64*) =10.14, p=0.002,η_p_^2^= 0.14; Figure 2A]. Bonferroni corrected posthoc testing revealed that while control male and female mice spent similar times in the central square of the open field (M: 34.06sec ± 6.05, F: 33.41 sec ± 5.39; *t*=0.39, p=1.00), among physically stressed mice, the time spent by female mice (9.65 sec ± 3.71) in the central square of the open field test was significantly less than the male mice (17.77sec ± 3.70; *t*=4.90, p<0.001). Between the different experimental conditions, control male mice spent a significantly higher time in the central square compared to the physically stressed male mice (*t*=9.83, p<0.001). Likewise control female mice spent a much higher time in the central square compared to physically stressed female mice (*t*=14.33, p<0.001).

**Figure 2.**
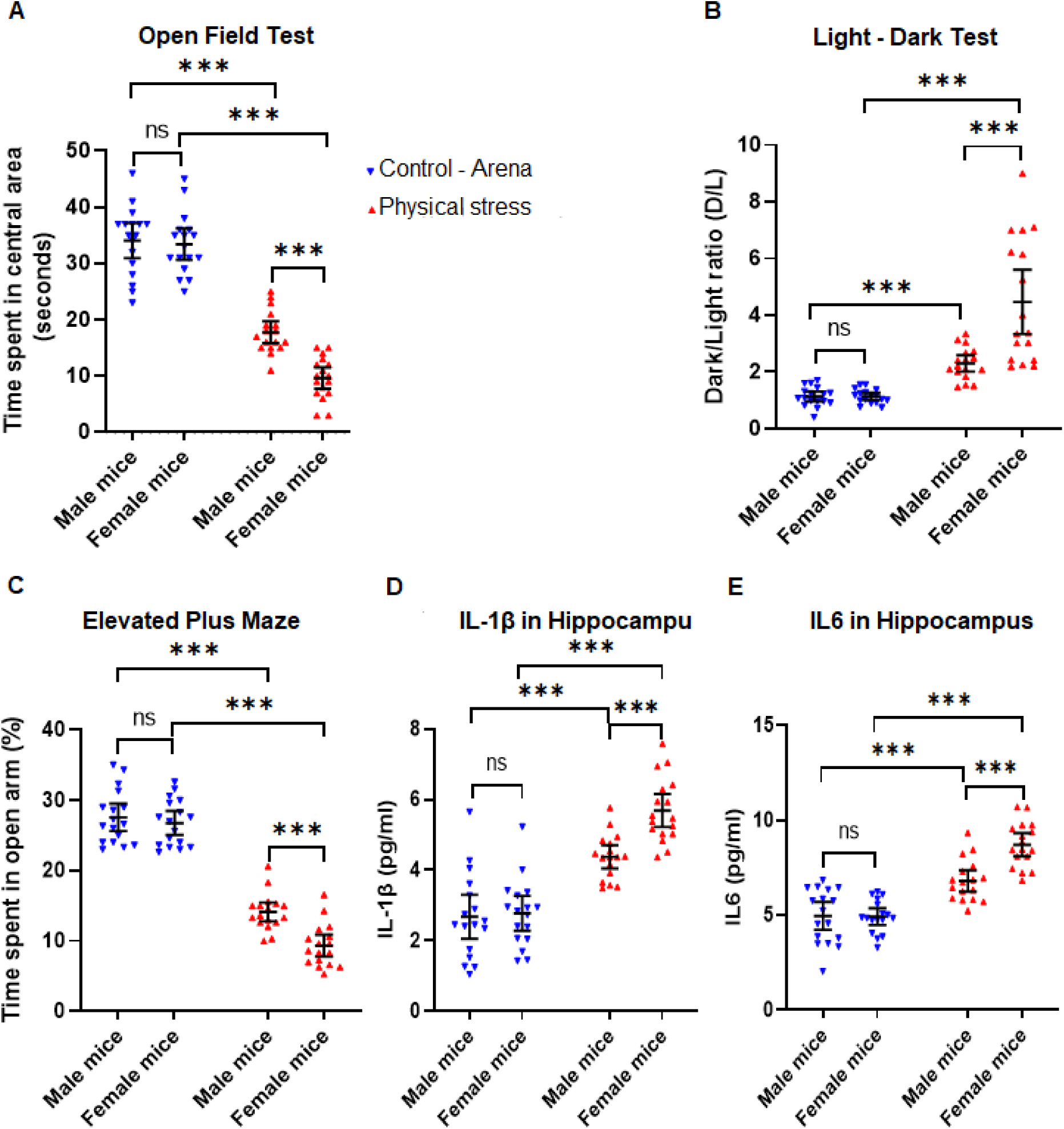
Female mice display an exacerbated stress response compared to male mice. **(A)** Open field test showing physically stressed mice exhibited significantly higher anxiety levels when compared to the controls, as illustrated by their reluctance to explore or spend more time in the central area of the OFT, with the female stressed mice showing greater anxiety. **(B)** Dark-Light test revealed female mice exposed to physical stress exhibited significantly higher anxiety levels (higher dark/light ratios) when compared to other groups, as explained by their hesitation to spend more time in the light chamber of the LDT. **(C)** Elevated plus maze revealed female mice subjected to physical stress displayed significantly increased anxiety levels compared to those in stressed males, as evidenced by their hesitation to spend more time in the open arms of the EPM. **(D)** Interleukin 1β (IL1β) levels in the hippocampus revealed stressed female mice showed aberrantly higher IL1β activity compared to those in stressed male mice. **(E)** Female mice subjected to physical stress showed aberrantly higher IL6 levels in the hippocampus relative to their male counterparts. All data are presented as Mean with 95% CI (n=17/group); ***p<0.001, ns (not significant); two-way ANOVA (Shapiro-Wilk’s, *p*>0.05) followed by Bonferroni *post-hoc* test.

In the light-dark test, analysis between subjects using factorial ANOVA on the D/L ratio revealed a significant main effect of stress [F (*1,64*) =64.51, p<0.001,η_p_^2^= 0.50], a significant main effect of sex [F (*1,64*) =15.2, p<0.001,η_p_^2^= 0.19] and a significant interaction between stress and sex [[F (*1,64*) =15.00, p<0.001,η_p_^2^= 0.19; Figure 2B]. Bonferroni post-hoc analysis showed that while control male and female mice had similar D/L ratios (M: 1.13 ± 0.34, F: 1.14 ± 0.24; *t*=0.001, p=1.00), physically stressed female mice had significantly higher D/L ratios compared to physically stressed male mice (M: 2.3 ± 0.56, F: 4.47 ± 2.19; *t*=5.48, p<0.001). Control male mice exhibited a significantly lower D/L ratio than physically stressed male mice (*t*=2.94, p=0.027), as did control female mice compared to physically stressed female mice (*t*=8.42, p<0.001).

Results from the elevated plus maze (time spent in the open arm of the maze) was also compared fully between-subjects using factorial ANOVA, and demonstrated a significant main effect of stress [F (*1,64*) = 390.60, p<0.001, η_p_^2^= 0.86], a significant main effect of sex,[F (*1,64*) = 12.85, p<0.001,η_p_^2^= 0.17], and a significant interaction between stress and sex,[F (*1,64*) = 6.519, p=0.013, η_p_^2^= 0.09; Figure 2C]. Once again, control male and female mice exhibited a similar amount of time spent in the open arm (M: 27.56 sec ± 3.295, F: 26.76 sec ± 3.82; *t*=0.88, p=1.00), but physically stressed female mice spent a significantly lower time in the open arm than physically stressed male mice (M: 14.12 sec ± 2.61, F: 9.33 sec ± 3.03; *t*=4.34, p<0.001). Within the different experimental conditions, the control male mice spent a significantly higher time in the open arm compared to the physically stressed mice (*t*=12.17, p<0.001) as did the female control mice compared to the physically stressed female mice (*t*=15.78, p<0.001). Thus far, in all three behavior assessments, stressed female mice displayed a hyper anxious behavior. Next, we wanted to investigate pro-inflammatory cytokines in the brain and see if they were consistent with the exaggerated anxiety response in the mice. To this end, we performed ELISA assays to test for hippocampal IL1β and IL6 in this cohort. In each case we analyzed the results with individual 2(arena control, physical stress) X 2(male, female) factorial ANOVAs followed with Bonferroni corrected pairwise comparisons for the significant main effects and interactions. For hippocampal IL1β, the factorial ANOVA revealed an extremely significant main effect of stress, [F (*1,64*) = 99.24, p<0.001, η_p_^2^= 0.61], a significant main effect of sex [F (*1,64*) = 9.32, p=0.003, η_p_^2^= 0.13] and a small yet significant interaction between stress and sex [F (*1,64*) = 9.32, p=0.011, η_p_^2^= 0.09; Figure 2D]. Consistent with the behavioral data, the stressed female mice had a significantly higher level of IL1β compared to the stressed male mice (M: 4.37 pg/ml ± 0.64, F: 5.69 pg/ml ± 0.91; *t*=4.02, p<0.001), while control male and female mice had similar levels of hippocampal IL1β (M: 2.68 pg/ml ± 1.21, F: 2.78 pg/ml ± 0.96; *t*=0.29, p=1.00). Within the different experimental conditions, physically stressed female mice had much higher levels than control female mice (*t*=8.91, p<0.001) and a similar pattern of difference was also observed between control males and stressed males, although the magnitude of the difference was less between control and stressed groups in the males (*t*=5.18, p<0.001).

For IL6, the factorial ANOVA revealed an extremely significant main effect of stress [F (*1,64*) = 100.79, p<0.001, η_p_^2^= 0.61], a significant effect of sex [F (*1,64*) = 11.074, p=0.001,η_p_^2^= 0.15] and a significant interaction [F (*1,64*) = 11.93, p<0.001, η_p_^2^= 0.16; Figure 2E]. Just like IL1β levels, hippocampal IL6 levels (pg/ml) were higher in stressed females compared to stressed males (M: 6.81 ± 1.10, F: 8.71 ± 1.17; *t*=4.80, p<0.001) but not in control males versus females (M: 4.96 ± 1.43, F: 4.93 ± 0.86; *t*=0.09, p=1.00). Stressed male and well as female mice displayed much higher IL6 in the hippocampus than control males and females (*t*=4.66, p<0.001; *t*=9.451, p<0.001, respectively). Therefore, we discovered a pattern of pro-inflammatory cytokine expression that mirrors the sex difference seen in the behavioral data following physical stress, compared to control animals. We repeated these experiments with animals exposed to predator odor stress and observed similar anxiety responses with the three above mentioned behavioral tests as well as elevations in hippocampal IL1β and IL6, as measured with ELISA (Supplementary Figure S1A-E). The sex differences were apparent in that cohort as well, however, the magnitude of the effects was lesser than what was elicited by the physical stress model. Keeping in mind the robust responses we saw following physical stress, hereafter, for all mechanistic investigation, we used the physical stress model and mice near stress experiment arena without a direct line of sight to the stress experiments as controls.

### Signaling pathways that lead to production of IL1β are upregulated in the hippocampus and amygdala of physically stressed female mice to a greater extent than male mice

Next, we focused on molecules that are involved in the production of IL1β in two brain areas which always figure prominently in the literature related to stress and anxiety-hippocampus and amygdala. Using immunoblot we probed these brain areas in physically stressed mice as well as control mice at one-week post-stress. We observed a robust induction of the NLRP3 inflammasome, as well as cleaved Caspase 1 (p20), the key molecules responsible for the generation of IL1β.

Consistent with the results shown in Figure 2, the induction NLRP3 and cleaved Caspase 1 (p20) were more prominent in stressed female mice, in comparison with stressed male mice and control group of mice, in both the hippocampus (Figure 3A) and the amygdala (Figure 3C). We normalized the NLRP3immunoblot band intensities against Tubulin and compared the relative band intensities from all groups with 2(control, physical stress) X 2 (male, female) factorial ANOVAs for hippocampus and amygdala each. For samples from hippocampus, the factorial ANOVA demonstrated a significant main effect for stress [F (*1,20*) = 84.89, p<0.001, η_p_^2^= 0.81], a significant main effect of sex [F (*1,20*) = 15.60, p<0.001, η_p_^2^= 0.44] on relative band intensities for NLRP3,qualified by significant interaction between stress and sex [F (*1,20*) = 12.90, p=0.002, η_p_^2^= 0.39; Figure 3B]. The stressed female mice had a significantly higher relative NLRP3 band intensity compared to the stressed male mice (M: 1.95 ± 0.5, F: 3.23 ± 0.61; *t*=5.33, p<0.001), while control male and female mice had similar levels of hippocampal NLRP3 (M: 1.0 ± 0.17, F: 1.06 ± 0.2; *t*=0.25, p=1.00). Within the different experimental conditions, physically stressed female mice had much higher NLRP3 levels than control female mice (*t*=9. 4, p<0.001) and a similar pattern of difference was also observed between control males and stressed males, although the magnitude of the difference was less between control and stressed groups in the males (*t*=3.96, p=0.005).

**Figure 3.**
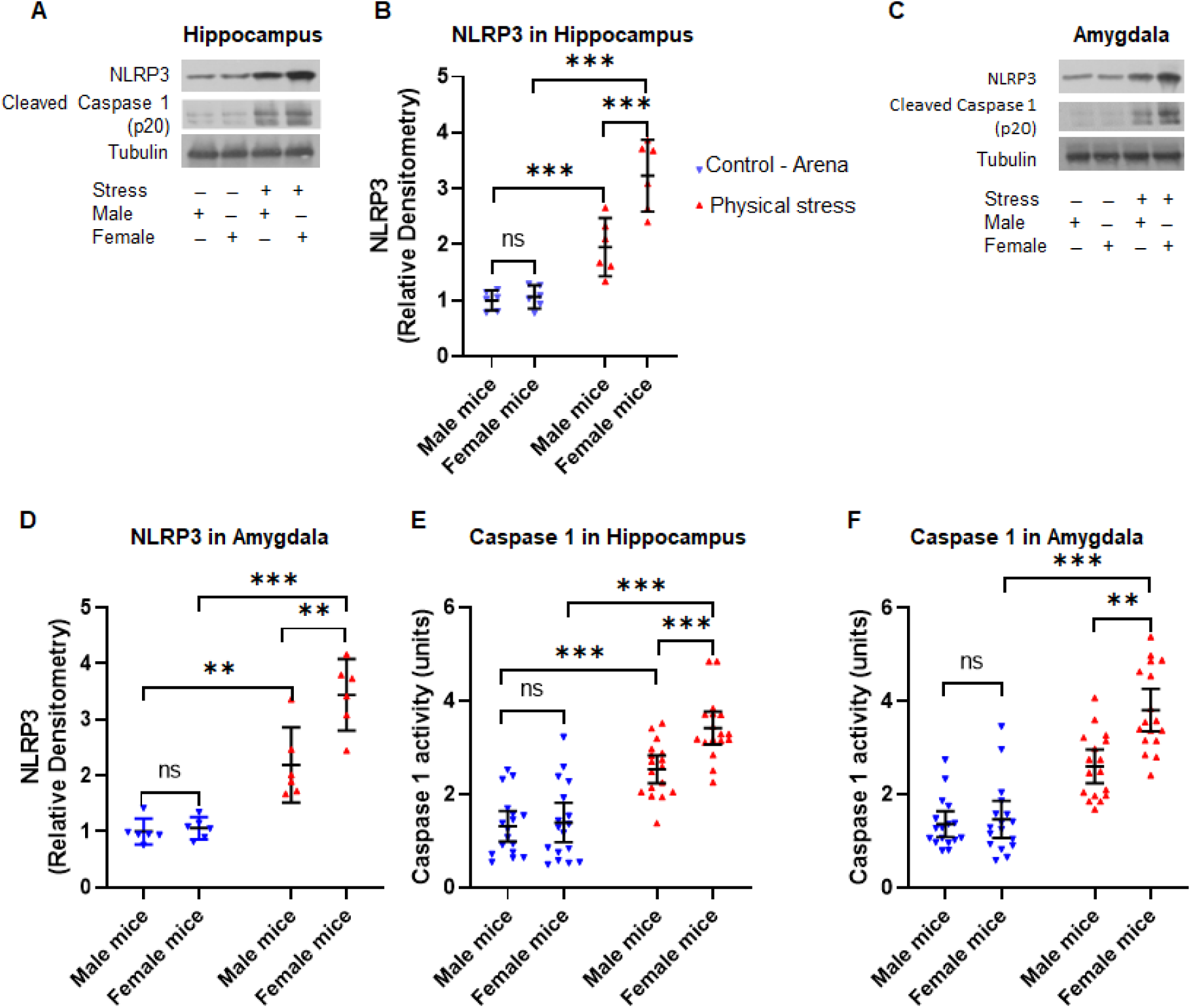
Physical stress leads to induction of NLRP3 inflammasome and Caspase 1 activation in hippocampus and amygdala of mice. **(A)** Immunoblot analysis of NLRP3 and cleaved Caspase 1 from hippocampus homogenates. **(B)** The relative densitometry of NLRP3 immunoblots of samples from the hippocampus, which shows physical stress-induced higher levels of NLRP3 inflammasome in female mice in comparison to male mice.**(C)** Western blot analysis of NLRP3 and cleaved Caspase 1 in amygdala homogenates. **(D)** The relative densitometry analysis of NLRP3 immunoblots of amygdala samples. The analysis showed stressed female mice exhibited increased induction of NLRP3 inflammasome in comparison to male mice. Caspase 1 activity in the hippocampus **(E)** and amygdala **(F)**, as measured in respective brain homogenates using Caspase 1 assay kit. Female mice subjected to physical stress showed significantly higher Caspase 1 activation relative to their male counterparts and control mice. All values are presented as Mean with 95% CI (n=6-17/group); ***p<0.001, ns (not significant); two-way ANOVA (Shapiro-Wilk’s, *p*>0.05) followed by Bonferroni *post-hoc* test.

The relative band intensities of immunoblot from amygdala samples also followed a similar pattern (Figure 3D). The factorial ANOVA demonstrated a significant main effect for stress [F (*1,20*) = 84.74, p<0.001, η_p_^2^= 0.82], a significant main effect of sex [F (*1,20*) = 11.91, p<0.001, η_p_^2^= 0.37] on relative band intensities for NLRP3from amygdala, qualified by significant interaction between stress and sex [F (*1,20*) = 9.87, p=0.005, η_p_^2^= 0.33; Figure 3D]. Once again the amygdala samples from stressed female mice had a significantly higher relative NLRP3 intensity as compared to the stressed male mice (M: 2.19 ± 0.64, F: 3.44 ± 0.61; *t*=4.66, p<0.001). Within the different experimental conditions, physically stressed female mice had significantly higher NLRP3 levels in amygdala than control female mice (*t*=8.88, p<0.001) and a similar pattern of difference was also observed between control males and stressed males, although the magnitude of the difference was less between control and stressed groups in the males (*t*=4.44, p=0.002). We also observed similar pattern from analysis of Caspase 1 activity in hippocampus (Figure 3E) and amygdala (Figure 3F). The physically stressed female mice had higher Caspase 1 activity as compared to their stressed male counterparts and controls. Similarly, the quantification of cleaved Caspase 1 relative band intensities from immunoblots showed significantly higher band intensities in physically stressed females as compared to stressed males and control groups in both amygdala (Supplementary Figure S2A) and hippocampus (Supplementary Figure S2B).

NLRP3 inflammasome play a critical role in the pathophysiology of neuroinflammation through activation of Caspase 1 and IL1β cytokine. Our data until here suggests that physical stress elicits hyper anxious behavior, robust NLRP3 induction in the hippocampus and amygdala, alongside activation of downstream mediators IL1β and Caspase 1, and all responses are more pronounced in females compared to males.

### Pharmacological inhibition of the NLRP3 inflammasome with MCC950 attenuates sexually divergent anxious behavior in mice as well as IL1β and Caspase 1 activity in the hippocampus and amygdala

Next, we carried out a pharmacological NLRP3 inhibition study, which served as proof-of-concept experiments to verify whether targeting NLRP3 pharmacologically can suppress the IL1β production and rescue the anxious behavior in mice subjected to restrained stress and underwater trauma. To this end, we selected MCC950, a compound known to inhibit NLRP3 and have an excellent blood-brain barrier permeability (Coll et al., 2015; Zahid et al., 2019). We injected control as well as physically stressed mice with vehicle and MCC950 (50mg/kg, i.p.) and tested the mice in the three behavioral paradigms mentioned before as well as analyzed their brains biochemically for IL1β and related markers. For each outcome measure we used 2(control, stressed) X 2(male, female) X 2 (vehicle, MCC950) Factorial ANOVAs. Given the pattern of interactions noted in the earlier set of experiments, we hypothesized *apriori* that we will see significant interactions among the factors, and the patterns of the interactions were especially of interest to us. We also wanted to see if the drug by itself influenced behavior and neuroinflammatory correlates in control mice, in addition to the drug’s ability to rescue phenotypes in stressed mice, as well as gender divergent effects.

When tested on the open field with the time spent in the central square of OFT and analyzed by a factorial ANOVA (Figure 4A), there was a significant main effect of stress [F (*1,168*) = 205.44, p<0.001, η_p_^2^= 0.55], a main effect of sex [F (*1,168*) = 4.68, p=0.032,η_p_^2^= 0.03], and a significant main effect of drug [F (*1,168*) = 52.86, p<0.001, η_p_^2^= 0.23]. Two interaction terms were significant, namely stress*sex [F (*1,168*) = 14.41, p<0.001, η_p_^2^= 0.08] and stress*drug [F (*1,168*) = 27.75, p<0.001, η_p_^2^= 0.14]. Control mice both vehicle treated (M: 30.35sec ± 6.04, F: 32.95 sec ± 8.69; *t*=0.66, p=1.00) or MCC950 treated (M: 33.31 sec ± 7.31, F: 34.05 sec ± 8.48; *t*=0.39, p=1.00) did not exhibit significant difference within sex or treatment (M: t=0.492,p=0.95). Vehicle treated mice subjected to physical stress, however, displayed a sharp decline in the time spent in the central square of the open field which with the decline being sharper in female mice than males (M: 16.64 sec ± 4.81, F: 8.05 sec ± 3.14) compared to control animals, both vehicle treated (*t*=13.86, p<0.001) as well as MCC950 treated (*t*=13.86, p<0.001). Physically stressed mice dosed with MCC950 showed a significant rescue from vehicle treated stressed mice (M: 26.32 sec ± 5.32, F: 23.14sec ± 6.52) compared to vehicle treated stressed mice (*t*=8.87, p<0.001).

**Figure 4.**
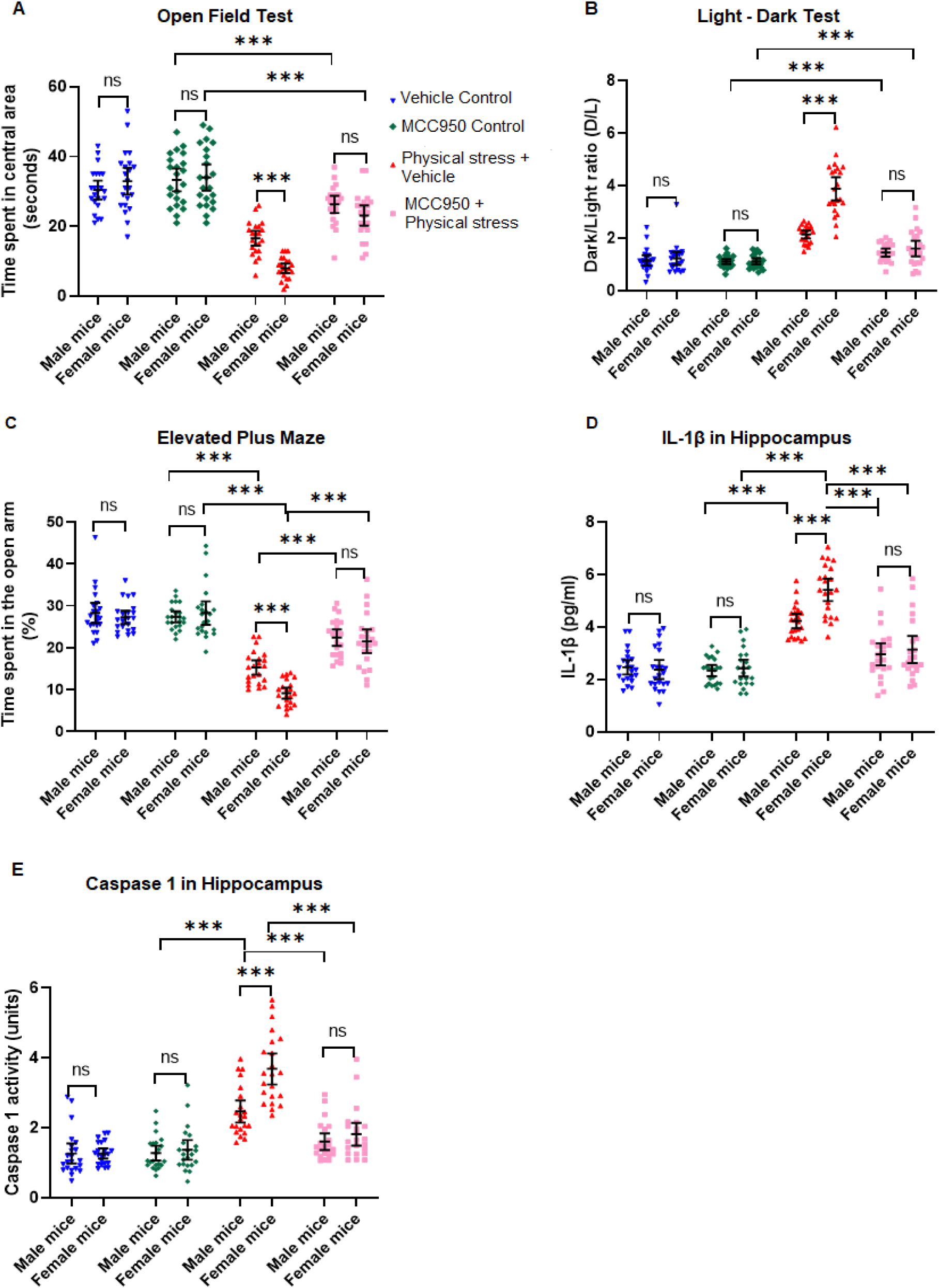
Inhibition of NLRP3 inflammasome by MCC950 treatment attenuates anxious behavior and neuroinflammation in stressed mice. Physical stress was induced by subjecting mice to restraint and underwater trauma. Control mice were treated with either vehicle or MCC950. Similarly, stressed mice were treated with either vehicle or MCC950. **(A)** Open Field Test: as compared to controls, physically stressed mice dosed with MCC950 exhibited significantly decreased anxiety levels, as illustrated by the increased time spent in the central area of the OFT, in both male and female mice. **(B)** Light-Dart Test: physically stressed mice dosed with MCC950 exhibited significant improvement from hyper-anxious behavior, as explained by the reduced D/L ratio, i.e. increased exploration time in the light chamber of the LDT. **(C)** Elevated plus maze test revealed physical stress mice dosed with MCC950 displayed significant rescue from anxiety compared to the vehicle controls, as evidenced by the increased duration of time spent in the open arms of the EPM. **(D)** Examination of IL1β in hippocampal homogenates by ELISA revealed physically stressed mice dosed with MCC950 showed diminished IL1β levels compared to the vehicle controls, denoting significant rescue from anxiety. **(E)** Caspase 1 activity in the hippocampus: physically stressed mice administered with MCC950 exhibited attenuated Caspase 1 activation relative to their vehicle controls. All values are presented as Mean with 95% CI (n=22/group); ***p<0.001, ns (not significant); two-way ANOVA (Shapiro-Wilk’s, *p*>0.05) followed by Bonferroni *post-hoc* test.

The light-dark test showed a similar pattern (Figure 4B). The factorial ANOVA conducted on the D/L ratios revealed a significant main effect of stress [F (*1,168*) = 191.405, p<0.001, η_p_^2^= 0.53], a main effect of sex [F (*1,168*) = 36.97, p<0.001, η_p_^2^= 0.18], and a significant main effect of drug [F (*1,168*) = 93.43, p<0.001, η_p_^2^= 0.36]. This was further qualified by significant interaction terms, including a 3-way interaction between stress, sex and drug [F (*1,168*) = 22.058, p<0.001, η_p_^2^= 0.12]. Control mice both vehicle treated (M: 1.15 ± 0.45, F: 1.22 ± 0.55; *t*=0.02, p=1.00) or MCC950 treated (M: 1.1± 0.22, F: 1.11± 0.26; *t*=0.98, p=1.00) did not exhibit significant difference within sex or drug treatment (Males, *t*=0.29, p=1.00; Females, *t*=0.71, p=1.00). Vehicle treated mice subjected to physical stress, however, showed a markedly increased D/L ratio, with female ratios being significantly higher (M: 2.141± 0.34, F: 3.89 ± 0.99; *t*=10.75, p<0.001) compared to control animals, both vehicle treated (M: *t*=6.01, p<0.001;F: *t*=16.4, p<0.001) as well as MCC950 treated (M: *t*=6.39, p<0.001; M: *t*=17.11, p<0.001). The D/L ratio was restored in physically stressed mice treated with MCC950 and showed a significant rescue (M: 1.45 ± 35, F: 1.6 ± 0.68, *t*=0.93, p=1.00) and their performance was not significantly different from MCC950 treated control mice (M: *t*=0.4, p=0.957; F: *t*=3.04, p=0.078).

Next, we analyzed the data from elevated plus maze (Figure 4C). We used the time spent by the mice from all groups, in the open arms, and calculated percentage time spent in the open arm. The factorial ANOVA conducted on the EPM data revealed a significant main effect of stress [F (*1,168*) = 230.08, p<0.001, η_p_^2^= 0.58], a main effect of sex [F (*1,168*) = 6.38, p=0.012, η_p_^2^= 0.04], and a significant main effect of drug [F (*1,168*) = 48.32, p<0.001, η_p_^2^= 0.22]. This was further qualified by significant interaction terms, with the most significant interaction between stress and drug [F (*1,168*) = 47.29, p<0.001, η_p_^2^= 0.22]. Control unstressed mice, both vehicle treated (M: 28.26± 5.47, F: 27.24 ± 3.59; *t*=0.72, p=1.00) or MCC950 treated (M: 27.35± 3.1, F: 28.36± 6.29; *t*=0.65, p=1.00) spent comparable times in the open arms and did not exhibit significant difference within sex or drug treatment (*t*=0.52, p=1.00). Vehicle treated mice subjected to physical stress, however, showed a markedly reduced percentage time spent in the open arm, with females spending significantly lesser time in the open arm of the EPM (M: 15.27± 3.9, F: 9.12 ± 2.96; *t*=6.15, p<0.001) compared to control animals, both vehicle treated (*t*=15.59, p<0.001) as well as MCC950 treated (M: *t*=15.64, p<0.001). The percentage time spent in the open arm was markedly improved in the physically stressed mice treated with MCC950 (M: 22.39± 4.4, F: 21.51 ± 6.37, *t*=0.62, p=1.00).

Thus far, the behavioral data replicated our earlier findings from Figure 2 with respect to all three behavioral paradigms used to measure anxious behavior. Our model of physical stress produced robust anxious behavior in mice, which was significantly more pronounced in female mice. When we inhibited the NLRP3 inflammasome, an important upstream molecule that mediates the activation of Caspase 1 and IL1β, using pharmacological inhibitor MCC950, mice showed significant rescue from heightened anxious behavior in all three behavior tests. Thus it implicates NLRP3-Caspase 1-IL1β pathway in anxiogenesis following physical stress.

We then turned to investigating biochemical changes in the brain following NLRP3 inhibition by administering MCC950. We measured hippocampal IL1β levels with ELISA and analyzed the data with a factorial ANOVA (Figure 4D).The factorial ANOVA conducted revealed a significant main effect of stress [F (*1,168*) = 153.65, p<0.001, η_p_^2^= 0.48], a main effect of sex [F (*1,168*) = 7.77, p=0.006, η_p_^2^= 0.04], and a significant main effect of drug [F (*1,168*) = 58.62, p<0.001, η_p_^2^= 0.24]. This was further qualified by significant interaction terms, with the most significant interaction between stress and drug [F (*1,168*) = 49.24, p<0.001, η_p_^2^= 0.23]. Control mice both vehicle treated (M: 2.47 pg/ml ± 0.62, F: 2.39 pg/ml ± 0.82; *t*=0.35, p=1.00) or MCC950 treated (M: 2.34 pg/ml ± 0.48, F: 2.44 pg/ml ± 0.72; *t*=0.65, p=1.00) had similar levels of IL1β in the hippocampus, did not show a sex difference or an upregulation in IL1β production just by the drug treatment(M: *t*=0.52, p=1.00; F:*t*=0.22, p=1.00). Vehicle treated mice subjected to physical stress, had significantly elevated hippocampal IL1β levels, with more prominent increase in females (M: 4.24 pg/ml ± 0.62, F: 5.42 pg/ml ± 0.62; *t*=4.79, p<0.001) compared to control animals, both vehicle treated (M: *t*=7.14, p<0.001;F: *t*=12.27, p<0.001) as well as MCC950 treated (M: *t*=7.66, p<0.001;F: *t*=12.06, p<0.001). When dosed with MCC950, IL-1β levels went down significantly, and abolished the difference between males and females, although females tended to have slightly elevated IL-1β even in treated animals (M: 2.97 pg/ml ± 0.96, F: 3.15 pg/ml ± 1.16, *t*=0.74, p=1.00). Following MCC950 treatment, hippocampal IL1β levels in physically stressed animals were not significantly higher than vehicle treated (M: *t*=1.99, p=1.00; F: *t*=3.08, p=0.07) or MCC950 treated (M: *t*=2.52, p=0.36; F: *t*=2.86, p=0.13) control mice.

We also analyzed the activity of Caspase 1 in the hippocampus, to validate our results in this pharmacological NLRP3 inhibition experiment, and analyzed the data with a Factorial ANOVA. Overall, the pattern of Caspase 1 activity data (Figure 4E) mirrored the IL1β data closely. Caspase 1 activity rose sharply and significantly in stressed mice treated with vehicle, and was significantly attenuated in physically stressed mice treated with NLRP3 inhibitor MCC950. We confirmed similar patterns of inhibition of IL1β as well as Caspase 1 activity in the amygdala of stressed mice treated with MCC950 (Supplementary Figure S3 A-B). Therefore, this experiment provided evidence that the NLRP3 inflammasome activation is a crucial part of the neuroinflammatory response seen in this model of physical stress and anxiogenic behavior, with females displaying a more severe phenotype compared to males.

### Bruton’s Tyrosine Kinase (BTK) is induced in hippocampus and amygdala of mice following physical stress in a sexually divergent pattern

To investigate a potential therapeutic avenue, we probed further upstream of NLRP3, and focused on BTK activation, a molecular event that is critical to the assembly and functioning of NLRP3 inflammasome. Phosphorylation of BTK at Tyr223 within the SH3 domain is necessary for the full activation of BTK and its downstream signaling, including induction of NLRP3 inflammasome. We subjected lysates from the hippocampus and amygdala of stressed and control mice to immunoblot analysis, using anti-phospho-BTK antibody (Tyr223, D9T6H, rabbit monoclonal antibody, 1:1000, #87141, Cell Signaling Technology),anti-BTK antibody (D3H5, rabbit monoclonal antibody, 1:1000, #8547, Cell Signaling Technology) and anti-alpha tubulin for loading control (ab4074, rabbit polyclonal antibody, 1:5000, Abcam). On conducting a 2X2 ANOVA (stress, sex) of tubulin-normalized pBTK relative densitometry from the hippocampal lysates revealed a significant main effect of stress[F (*1,20*) =110.63, p<0.001,η_p_^2^= 0.85]and a significant main effect of sex [F (*1,20*) =13.03, p=0.002, η_p_^2^= 0.4]. This was further qualified by a significant interaction between the two factors [F (*1,20*) =14.97, p<0.001, η_p_^2^= 0.43; Figure 5A-B]. Bonferroni corrected post-hoc testing revealed that physical stress induced an approximately 2 and 3 fold increase in phospho-BTK in male and female mice, respectively (M: 2.058± 0.44, F: 3.25 ± 0.56; *t*=5.29, p<0.001), compared to control male and female mice (M: 1.00 ± 0.16, F: 0.96 ± 0.28; *t*=998, p=1.00).Between the two experimental conditions the upregulation in phopspho-BTK was much starker in stressed females as compared to control female mice (*t*=10.17, p<0.001) than stressed males and control male mice (*t*=4.70, p=0.002), although both sexes demonstrated a significant upshot of phospho-BTK following stress. There was no change in total BTK levels in all the groups, as revealed by immunoblots probed with BTK antibody (Figure 5A). Very similar patterns and effect sizes were found in the amygdala samples (Figure 5C-D). Immunoblot analysis of amygdala samples revealed that phospho-BTK was upregulated following physical stress, but disproportionately more significantly in females than male mice. Thus far, these experiments provided the evidence that BTK, an upstream regular of NLRP3, is activated in hippocampus and amygdala of mice following physical stress in sexually divergent pattern.

**Figure 5.**
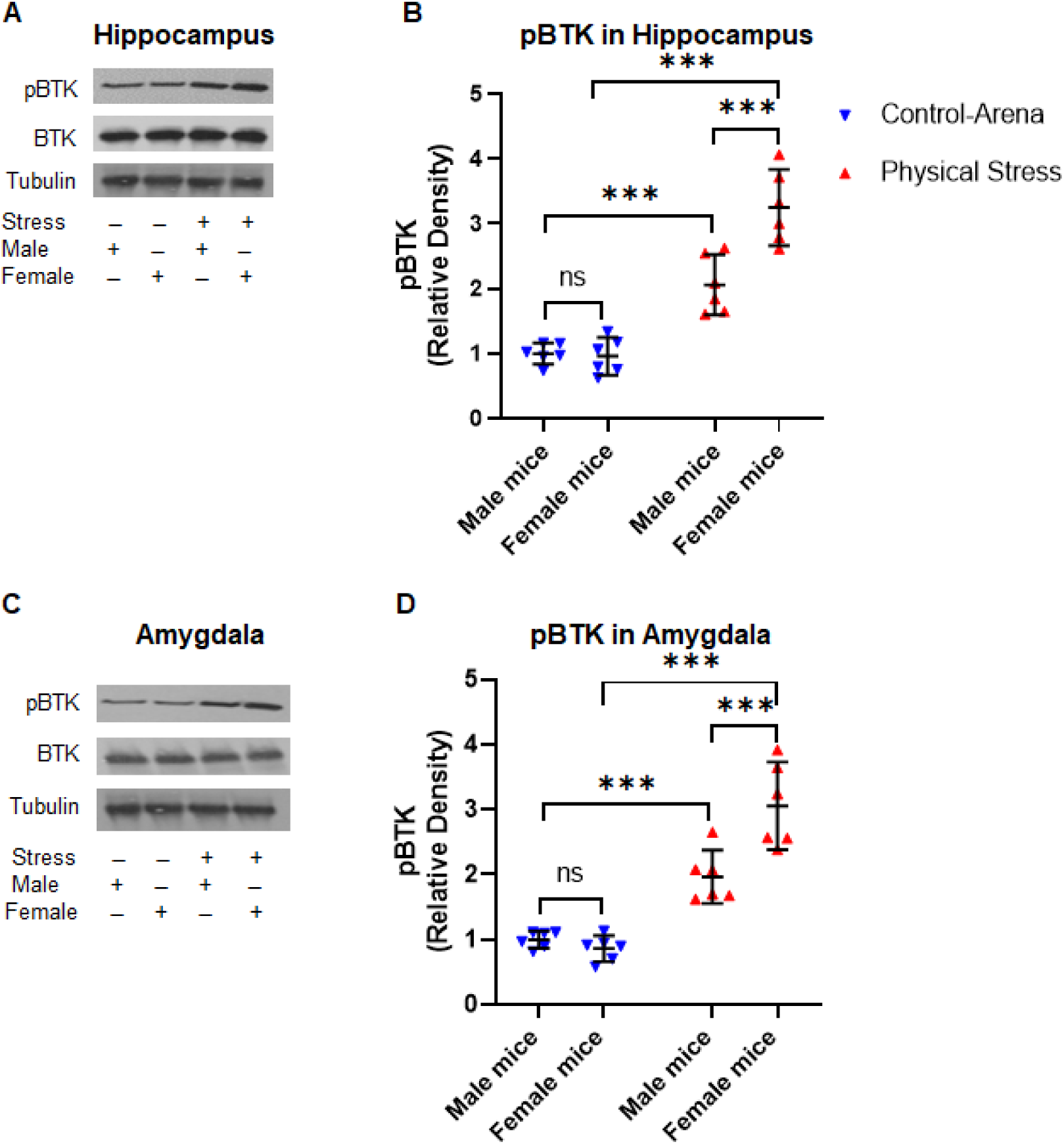
Physical stress in mice induces the activation of BTK in the hippocampus and amygdala. Mice were subjected to physical stress by restraint and underwater trauma. **(A)** Immunoblot analysis of the hippocampal homogenates using anti-phopsho-BTK (pBTK, Tyr223), BTK, and tubulin antibodies. **(B)** Relative densitometry analysis of pBTK immunoblots showing physical stress significantly induced the activation of BTK, as revealed by increased pBTK levels in stressed mice as compared to controls. **(C)** Western blot analysis of pBTK, total BTK, and tubulin as a loading control. **(D)** The relative densitometry analysis of pBTK immunoblots of amygdala samples. The analysis from both hippocampal and amygdala samples showed stressed female mice exhibited significantly increased induction of pBTK relative to their male counterparts and control mice. Data are presented as Mean with 95% CI (n=6/group); ***p<0.001, ns (not significant); one-way ANOVA (Shapiro-Wilk’s, *p*>0.05) followed by Bonferroni *post-hoc* test.

### Pharmacological inhibition of BTK produces anxiolysis and attenuates NLRP3, Caspase 1 and IL1β upregulation in hippocampus and amygdala of stressed mice

Several recent studies in different disease models, such as ischemic brain injury, cardiac dysfunction, and inflammatory diseases have suggested that BTK is an upstream positive-regulator of NLRP3 inflammasome, which in turn induce downstream proinflammatory molecular events such as activation of Caspase 1 and IL1β (Ito et. al. 2015, Liu et. al. 2017, O’Riordan et. al. 2019, Purvis et. al. 2020, Franke et. al. 2020, Bittner et. al. 2020). Utilizing Ibrutinib, as a FDA approved BTK inhibitor, these studies have suggested BTK as a therapeutically relevant NLRP3 regulator. Ibrutinib binds irreversibly to BTK and inhibits the phosphorylation of Tyr 223, thus blocking BTK activity. In a proof-of-concept study we asked if inhibiting BTK with Ibrutinib would provide protection in physical stress model. We injected Ibrutinib (3 mg/Kg, i.p.) into control groups of mice and as well as those subjected to restraint and underwater trauma, and assessed them on behavioral and molecular end points.

First, we wanted to see if Ibrutinib injected stressed animals demonstrated a change in the levels of NLRP3 inflammasome, a key mediator for downstream inflammatory cascade. To this end we isolated hippocampi from stressed animals treated with vehicle or Ibrutinib and subjected the hippocampal lysates to Western blot analysis (Figure 6A). We observed that Ibrutinib inhibited the induction of NLRP3 in physically stressed mice, as compared to control group of mice. We compared the relative density of NLRP3 normalized with tubulin and analyzed the groups by a one-way ANOVA test (Figure. 6B). The group means were significantly different [F (5,30)= 33.95, p <0.001, η^2^= 0.85]. Bonferroni corrected Post-hoc testing revealed significantly high upregulation of NLRP3 in stressed females as compared to their male counterparts (p<0.001). Further, the relative density of NLRP3 in Ibrutinib treated stressed male and female mice was significantly reduced as compared to their stressed counterparts administered with only vehicle control (p<0.001).

**Figure 6.**
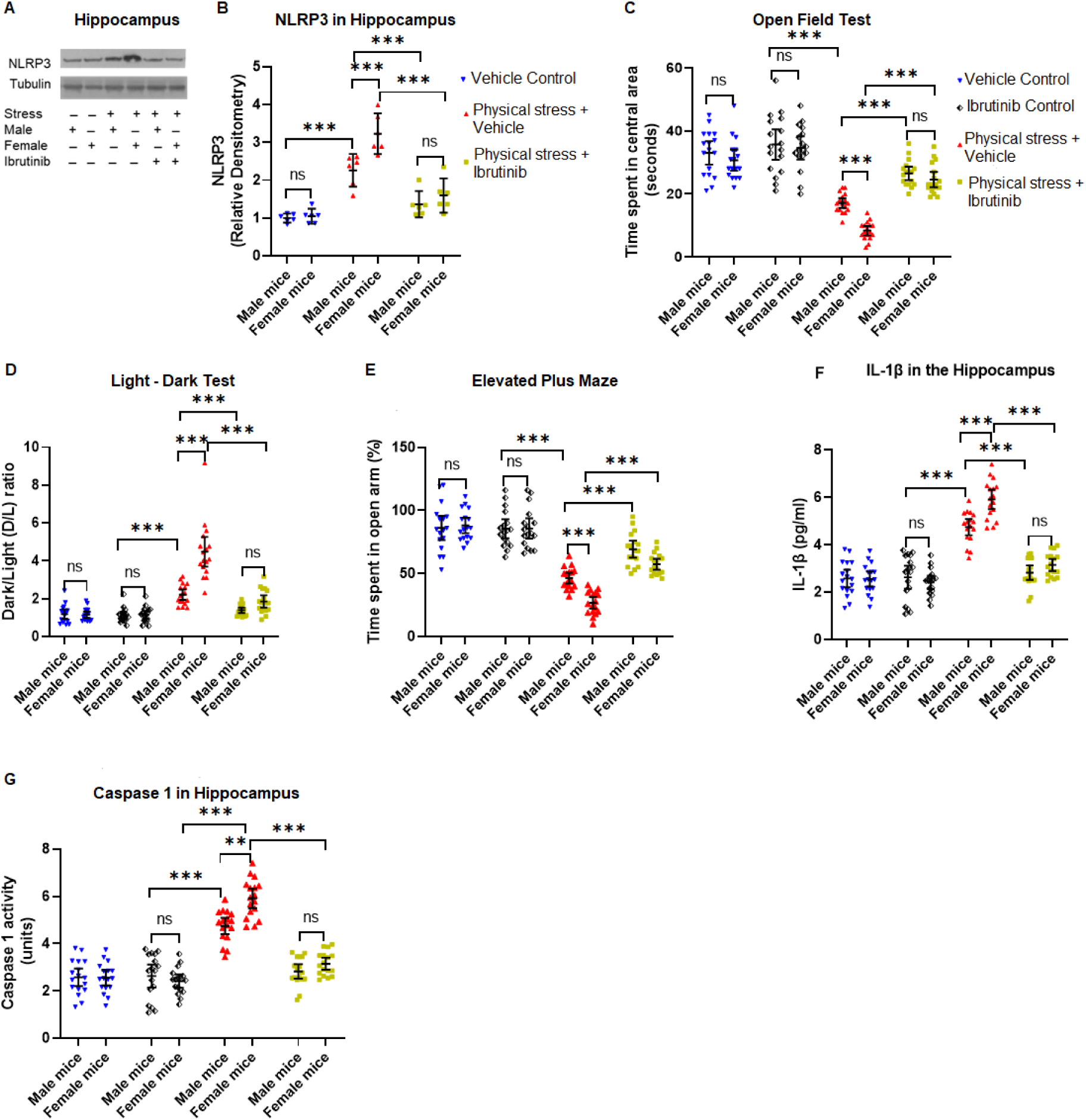
Pharmacological inhibition of BTK with Ibrutinib attenuates hyper-anxious behavior as well as pro-inflammatory molecules in the hippocampus of stressed mice. Mice subjected to restraint stress and underwater trauma were injected with Ibrutinib (3 mg/Kg, i.p.) or vehicle. **(A)** Immunoblot analysis of NLRP3 in hippocampus homogenates. **(B)** The relative densitometry of NLRP3 immunoblots from hippocampal samples showing Ibrutinib treatment significantly reduced the NLRP3 inflammasome levels in physically stressed mice as compared to vehicle-treated stressed mice. **(C)** Open field test: physically stressed mice dosed with Ibrutinib exhibited significantly decreased anxiety levels when compared to the vehicle controls, as illustrated by increased exploration time in the central area of the OFT.**(D)** Light-Dark test: physically stressed mice dosed with Ibrutinib exhibited significant rescue from hyper-anxiety as explained by the significant increase in their time spent in the light chamber of the LDT.**(E)** Elevated plus maze test: physically stress mice injected with Ibrutinib displayed significantly reduced anxiety, as evidenced by the increased duration of time spent in the open arms of the EPM.**(F)** Analysis of IL1β in the hippocampus by ELISA revealed physically stressed mice dosed with Ibrutinib showed diminished proinflammatory IL1β. **(G)** Caspase 1 activity in the hippocampus: administration of Ibrutinib in physically stressed mice showed reduced Caspase 1 activation, elucidating significant rescue from the anxiogenic proinflammatory pathway. All values are presented as Mean with 95% CI (n=22/group); ***p<0.001, ns (not significant); two-way ANOVA (Shapiro-Wilk’s, *p*>0.05) followed by Bonferroni *post-hoc* test.

Next, we carried out behavioral analysis on control and stressed animals and analyzed them by sex as well as drug treatment. For each experiment, we used separate 2(stressed, control) X 2 (male, female) X 2(vehicle, Ibrutinib) Factorial ANOVAs. We used same three behavioral tests described above. When the time spent in the central square of open field test was analyzed by a factorial ANOVA (Figure 6C), there was a significant main effect of stress [F (*1,128*) = 193. 19, p<0.001, η_p_^2^= 0.6], a main effect of sex [F (*1,128*) = 11.63, p<0.001, η_p_^2^= 0.08], and a significant main effect of drug [F (*1,128*) = 60.654, p<0.001, η_p_^2^= 0.32]. The interaction between stress and drug was significant with the largest effect size [F (*1,128*) = 21.202, p<0.001, η_p_^2^= 0.14]. Control mice both vehicle treated (M: 33.06 sec ± 7.24, F: 30.71sec ± 6.45; *t*=1.16, p=1.00) or Ibrutinib treated (M: 35.71 sec ± 9.38, F: 34.65 sec ± 7.19; *t*=0.51, p=1.00) did not exhibit significant difference within sex or treatment (M: *t*=1.27, p=1.00; F: *t*=1.92, p=1.00). Vehicle treated mice subjected to physical stress, however, displayed a sharp decline in the time spent in the central square of the open field, more prominently in the females (M: 17.12± 3.04, F: 8.35 ± 2.83; *t*=4.235, p<0.001) compared to control animals, both vehicle treated (M:*t*=7.703, p<0.001; F: *t*=10.8, p<0.001) as well as Ibrutinib treated (M: *t*=8.98, p<0.001; F: *t*=12.71, p<0.001). Ibrutinib treated mice subjected to physical stress showed significant rescue from hyper anxious behavior as compared to vehicle treated stressed mice in both males (M: 26.53 sec ± 4.16, *t*=4.548, p<0.001.) and females (F: 24.59 sec ± 4.77; *t*=7.85, p<0.001). After Ibrutinib treatment, physically stressed males and females spent similar time in the central square of the open field (*t*=0.94, p=1.00).

The light-dark test showed a similar pattern (Figure 6D) in Ibrutinib treated mice. The factorial ANOVA conducted on the D/L ratios revealed a significant main effect of stress [F (*1,128*) = 60.23, p<0.001, η_p_^2^= 0.50], a main effect of sex [F (*1,128*) = 33.98, p<0.001, η_p_^2^= 0.21], and a significant main effect of drug [F (*1,128*) = 56.62, p<0.001, η_p_^2^= 0.31]. This was further qualified by significant interaction terms, including a 3-way interaction between stress, sex and drug [F (*1,128*) = 15.81, p<0.001,η_p_^2^= 0.11]. Control mice both vehicle treated (M: 1.19± 0.48, F: 1.16 ± 0.33) or Ibrutinib treated (M: 1.12± 0.38, F: 1.15± 0.26) did not exhibit significant difference within sex or drug treatment. Vehicle treated mice subjected to physical stress, however, showed a markedly increased D/L ratio, with female ratios being significantly higher (M: 2.21± 0.54, F: 4.48 ± 1.53; *t*=9.68, p<0.001) compared to control unstressed animals, both vehicle treated (M: *t*=4.36, p<0.001;F: *t*=14.17, p<0.001) as well as Ibrutinib treated (M: *t*=3.49, p=0.02;F: *t*=14.22, p<0.001). The D/L ratio was significantly reduced in physically stressed mice treated with Ibrutinib, as compared to vehicle treated stressed mice (M: 1.5± 0.29, F: 1.86 ± 0.62, *t*=1.9, p=1.00). As compared to controls, Ibrutinib clearly improved the anxious behavior in stressed mice as evident by increased time spent in light chamber of the light-dark test (p<0.001).

Next, we analyzed the data from elevated plus maze test (Figure 6E). The factorial ANOVA conducted on the EPM data revealed a significant main effect of stress [F (*1,128*) = 268.113, p<0.001, η_p_^2^= 0.68], a significant main effect of sex [F (*1,128*) = 11.06, p=0.001, η_p_^2^= 0.08] and a significant main effect of drug [F (*1,128*) = 32.27, p<0.001, η_p_^2^= 0.2]. This was further qualified by significant interaction terms, with the most significant interaction between stress and drug [F (*1,128*) = 40.33, p<0.001, η_p_^2^= 0.24]. Control mice, both vehicle treated (M: 28.69± 6.21, F: 29.32 ± 4) or Ibrutinib treated (M: 28± 3.1, F: 28.36 ± 6.29) spent comparable times in the open arms and did not exhibit significant difference within sex or drug treatment (*t*=0.99, p=1.00). Vehicle treated mice subjected to physical stress, however, showed a markedly reduced time spent in the open arm, with females spending significantly lesser time in the open arm of the EPM (M: 15.44 ± 2.82, F: 8.9 ± 3.04) compared to control animals, both vehicle treated (*t*=16.07, p<0.001) as well as Ibrutinib treated (M: *t*=15.59, p<0.001). Treatment with Ibrutinib significantly reduced the anxiety in stressed mice, as evident by significant increase in exploration time in the open arm of EPM, as compared to controls (p<0.001). The percentage time spent in the open arm of EPM was markedly improved in both male and female stressed mice treated with Ibrutinib (M: 23.08± 4.33, F: 19.08 ± 2.81).

Thus far, our model of physical stress produced robust anxious behavior in mice, which was significantly more pronounced in female mice. When we inhibited BTK, an upstream regulator of NLRP3 inflammasome, with a pharmacological inhibitor, in all three behavior tests mice showed significant rescue from heightened anxious behavior, validating the implication of BTK pathway in anxiogenesis following physical stress.

We then turned to investigating biochemical changes in the brain following Ibrutinib administration. We measured hippocampal IL1β levels with ELISA and analyzed the data with a factorial ANOVA (Figure 6F). The factorial ANOVA conducted revealed a significant main effect of stress [F (*1,128*) = 153.65, p<0.001, η_p_^2^= 0.59], a main effect of sex [F (*1,128*) = 7.06, p=0.009, η_p_^2^= 0.05], and a significant main effect of drug [F (*1,128*) = 101.88, p<0.001, η_p_^2^= 0.44]. This was further qualified by significant interaction terms, with the most significant interaction between stress and drug [F (*1,128*) = 94.64, p<0.001, η_p_^2^= 0.43]. Control mice both vehicle treated (M: 2.58 pg/ml ± 0.73, F: 2.55 pg/ml ± 0.62) or Ibrutinib treated (M: 2.63 pg/ml ± 0.95, F: 2.41 pg/ml ± 0.55) had similar levels of IL1β in the hippocampus, did not show a sex difference or an upregulation in IL1β production just by the drug treatment (*t*=0.99, p=1.00). Vehicle treated mice subjected to physical stress, had significantly elevated hippocampal IL1β levels, with more prominent increase in females (M: 4.75 pg/ml ± 0.67, F: 3.15 pg/ml ± 0.49) compared to control animals, both vehicle treated (*t*=16.51, p<0.001) as well as Ibrutinib treated (*t*=14.02, p<0.001). When dosed with Ibrutinib, IL1β levels went down significantly in stressed mice, and abolished the difference between males and females, although females tended to still have slightly elevated IL1β in treated animals (M: 2.82 pg/ml ± 0.59, F: 3.15 pg/ml ± 0.5).

We also analyzed the activity of Caspase 1 in the hippocampus, to validate our results in this pharmacological inhibition experiment, and analyzed the data with a Factorial ANOVA. Overall, the pattern of Caspase 1 activity data (Figure 6G) mirrored the IL1β data closely. Caspase 1 activity rose sharply and significantly in stressed mice which were vehicle treated, however, Ibrutinib treatment significantly attenuated the Caspase 1 activity in hippocampus of both male and female stressed mice (p<0.001). Therefore, this experiment provided evidence that the BTK inhibition, thus likely reduction of NLRP3 inflammasome, is crucial part of the neuroinflammatory response seen in this model of physical stress and anxiogenic behavior, with females displaying a more severe phenotype compared to males. To further validate our observations with Ibrutinib treatment, we repeated this experiment with another compound, LFM-A13, also known in the literature for its ability to specifically suppress BTK (Ito et al. 2015). Control or stressed mice were injected with LFM-A13 (50 mg/Kg, i.p.) or vehicle and subjected to full battery of anxiety behavior tests and biochemical analysis. Our findings from LFM-A13 mirrored the findings from the Ibrutinib study (Supplementary Figure S4 A-E). Inhibition of BTK with LFM-A13 significantly suppressed the anxiety behavior in both male and female stressed mice, as compared to control groups (p<0.001). Further, LFM-A13 significantly reduced the activation of proinflammatory mediators Caspase 1 and IL1β in hippocampus (p<0.001). In total, our study suggested an important role of BTK in anxiogenic pathway.

## Discussion

The present study investigated the proinflammatory NLRP3 - Caspase 1 - IL-1β pathway, which recent research has indicated as a key mediator of heightened inflammatory response in several neuropathological conditions. We have used two different models of stress, first, a direct model of single prolonged immobilization stress, and forced swim followed by brief submersion underwater; second, predator odor which was an operationalized form of psychological stress. Both are established stress models in rodents (Verbitsky et al., 2020). For behavioral outcomes we used a set of well validated tests of anxiety in rodents (Walf and Frye, 2007; Crawley and Bailey, 2008; Solomonow and Tasker, 2015). To control for stress on account of relocation to stress behavior mouse facility, we consistently used control animals that were relocated from the animal facility and housed in the room where stress experiments were conducted, yet to prevent vicarious stress, they had no direct view or exposure to the arena or the mice undergoing stress experiments, precluding any olfactory or visual cues to stress.

Our study indicates that both models of stress used elicits a sharp increase in anxious behavior and neuroinflammatory mediators and that this increase in both behavior and inflammatory profile is significantly more pronounced in females compared to males. The presence of a pronounced neuroinflammatory response post-stress is well established in several preclinical rodent models and has been reviewed extensively (Mellon et al., 2018b; Enomoto and Kato, 2021). Elevation of a range of peripheral inflammatory mediators have been consistently observed in PTSD patients compared to age matched controls, both at protein and genetic levels (Passos et al., 2015; Calcia et al., 2016; Aliev et al., 2020; Zass et al., 2017; Yang and Jiang, 2020). The sexually dimorphic patterns of anxious behavior and the upregulation of neuroinflammatory markers we observed in our model are also consistent with three lines of evidence. First, in humans stress related disorders affect more women than men (Friedman et al., 2011; Bryant, 2019), as do anxiety related disorders (Donner and Lowry, 2013). Second, Lasselin and colleagues in their review of several human experimental stress models has suggested that the neuroinflammatory response in discussed models is overall, more robust in females than males (Lasselin et al., 2018). Finally, the sexually dimorphic neuroinflammatory response with females showing more robust neuroinflammation post-stress is noted in several rodent models of stress (Pyter et al., 2013; Bollinger et al., 2016; Fonken et al., 2018; Liu et al., 2019).

The NLRP3 inflammasome has been recently the target of intense research as a molecule of therapeutic interest in Alzheimer’s disease (Heneka et al., 2013; Dempsey et al., 2017), amyotrophic lateral sclerosis (Debye et al., 2018), vascular dementia (Lénárt et al., 2016), multiple sclerosis and experimental autoimmune encephalopathy (Gris et al., 2010; Inoue et al., 2012; Barclay and Shinohara, 2017), stroke (Fann et al., 2013; Espinosa-Garcia et al., 2020), bacterial meningitis (Hoegen et al., 2011; Geldhoff et al., 2013), traumatic brain injury (Ismael et al., 2018), cerebral hemorrhage (Feng et al., 2015), in addition to systemic illnesses such as peritoneal fibrosis (Hishida et al., 2019) and Type-2 diabetes (Masters et al., 2010), in both clinical and preclinical studies. The NLRP3 inflammasome has also received attention in the literature pertaining to psychiatric disorders (Frank et al., 2016; Kaufmann et al., 2017). The activation of the NLRP3 inflammasome has been observed in the mononuclear blood cells from patients with major depressive disorder (Alcocer-Gómez et al., 2014). Due to an established literature on the elevation of inflammatory markers following stress, as well as the documented role of the NLRP3 inflammasome activation in the maturation of IL1β, a powerful cytokine that has often been regarded as a “master regulator” of the neuroinflammatory response, the NLRP3 activation has also been proposed as a valuable candidate for therapeutic exploitation in stress related disorders (Alcocer-Gómez et al., 2017). Preclinical models of stress-induced depression, or foot-shock based stress models have utilized interfering with NLRP3 inflammasome activation using genetic knockout (Alcocer-Gómez et al., 2016; Su et al., 2017; Dong et al., 2020b) as well as pharmacological inhibition using several different pharmacological inhibitors (Zhang et al., 2020, 2015). The suppression of NLRP3 activation has led to alleviation of behavioral deficits in the said models. In that, our results are consistent with the existing literature. However, while we have validated the trends in literature in our work, there are several novel aspects of our work. We inhibited NLRP3 with MCC950, one of the most potent and specific small molecule inhibitors in the context of a single prolonged stress episode as well as psychological stress.

We have performed these experiments in both female and male mice, which are distinct from considerable bulk of research in this field, done exclusively on male animals. We have also provided a thorough characterization of molecular and behavioral end points. Especially, we clearly demonstrated the heightened activation of proinflammatory NLRP3 - Caspase 1 - IL1β pathway in the hippocampus and amygdala of stressed mice, which was significantly more pronounced in females as compared to their male counterparts. We also demonstrated an attenuated neuroinflammatory response in terms of IL1β, Caspase 1 activity concomitant to MCC950 based NLRP3 inhibition consistent with anxiolysis in both female and male animals.

We further probed upstream of the NLRP3 inflammasome and zeroed in on Bruton’s Tyrosine Kinase that acts as a positive regulator of NLRP3 activation (Ito et al., 2015a; Weber et al., 2017; Bittner et al., 2019). These studies demonstrated that BTK is required for NLRP3 mediated maturation of IL1β from pro-IL1β in macrophages. For the first time, our study provided evidence that stress induces the activation of BTK in both hippocampus and amygdala, further in a sexually dimorphic way, with female showing much more pronounced pBTK levels in brain. Several earlier studies made use of Ibrutinib, a selective inhibitor of BTK that is already FDA approved for oral clinical use in certain types of lymphomas and leukemia (De Claro et al., 2015; Wilson et al., 2015; Grommes et al., 2017; Lionakis et al., 2017; Kapoor and Ansell, 2018). Using a murine model of modified middle cerebral artery occlusion for 60 minutes followed by reperfusion, Ito and colleagues, demonstrated that Ibrutinib treatment for up to 12 hours post-reperfusion minimized infarct volume, improved neurological scores at recovery and markedly attenuated levels of the proinflammatory cytokines IL1β, IL6 and TNF-α (Ito et al., 2015b). In our study, we observed that Ibrutinib treated animals displayed an almost 40-50% rescue in anxious behaviors and a concomitant reduction in Caspase 1 activity and IL1β, in both hippocampus and amygdala, which was more pronounced in female animals. Since, we wanted to further validate our findings with BTK inhibition, we repeated the pharmacological inhibition experiment with LFM-A13, a second selective BTK inhibitor (Mahajan et al., 1999; Uckun et al., 2002; Bam et al., 2013) in the single prolonged stress model. The results closely mirrored that from the Ibrutinib experiment, further suggesting a potential role of BTK in modulating anxious behavior and proinflammatory pathway. In summary, we present novel and robust proof-of-concept findings, and provide the first evidence of anxiolysis and attenuated neuroinflammation in a model of single prolonged stress, following inhibition of BTK, an upstream regulator of NLRP3 inflammasome with Ibrutinib and LFM-A13. The effect of pharmacologic inhibition of BTK closely mirrors that of NLRP3 inhibition by MCC950, including behavioral rescue and down-regulation of Caspase 1 activity as well as levels of IL1β, suggesting that BTK inhibition interferes with NLRP3 activation and limits Caspase1 activity, ultimately lowering the levels of mature IL1β. This is also to our knowledge, the first *in-vivo* therapeutic application of LFM-A13 in a stress model, with clear effects on neuroinflammatory markers in hippocampus. Stress leads to an exacerbated neuroinflammatory response and significantly more anxiety in female mice, and with all three inhibitors viz. MCC950, Ibrutinib and LFM-A13, the rescue in female animals were also more pronounced than in males.

Ibrutinib, which is already FDA approved for use in certain types of cancers, can be explored further as a potential therapeutic molecule, as well as further development of LFM-A13 or other BTK inhibitors. Especially for Ibrutinib, a chemotherapeutic drug, side effect and tolerance data has been reviewed extensively (Roschewski et al., 2020). However, the drug has been administered in patients of central nervous system lymphoma (Lionakis et al., 2017), and recently in patients of severe COVID-19 to prevent the initiation of the cytokine storm (Roschewski et al., 2020), suggesting a careful titration of dosage under monitored condition can theoretically be conceived for other conditions. Because of its multiple activators as a pattern recognition receptor, directly inhibiting NLRP3 inflammasome assembly may interfere with protective inflammatory responses, making BTK a more suitable molecule of interest for targeting drug development.

Trauma and stress related disorders, comorbid with depression and anxiety is a major cause of debilitating psychiatric illness, which is already being indicated as a massive challenge due to the stressful situations globally created at the wake of the Covid-19 pandemic (Chew et al., 2020; Dubey et al., 2020). Urgent research is needed to find feasible therapeutic alternatives beyond the accepted SSRI-SNRI based treatment regimens for such disorders. In this context, further careful examination and testing of the therapeutic potential of Ibrutinib for use in such psychiatric illnesses can prove extremely beneficial.

## Acknowledgements

This study was supported by an intramural research grant to S.G. from Ashoka University, Brain Aneurysm Foundation and Ramalingaswamy Fellowship to I.S. and an Ashoka University graduate student fellowship to Z.M.

## Supplementary Figures

**Figure S1.**
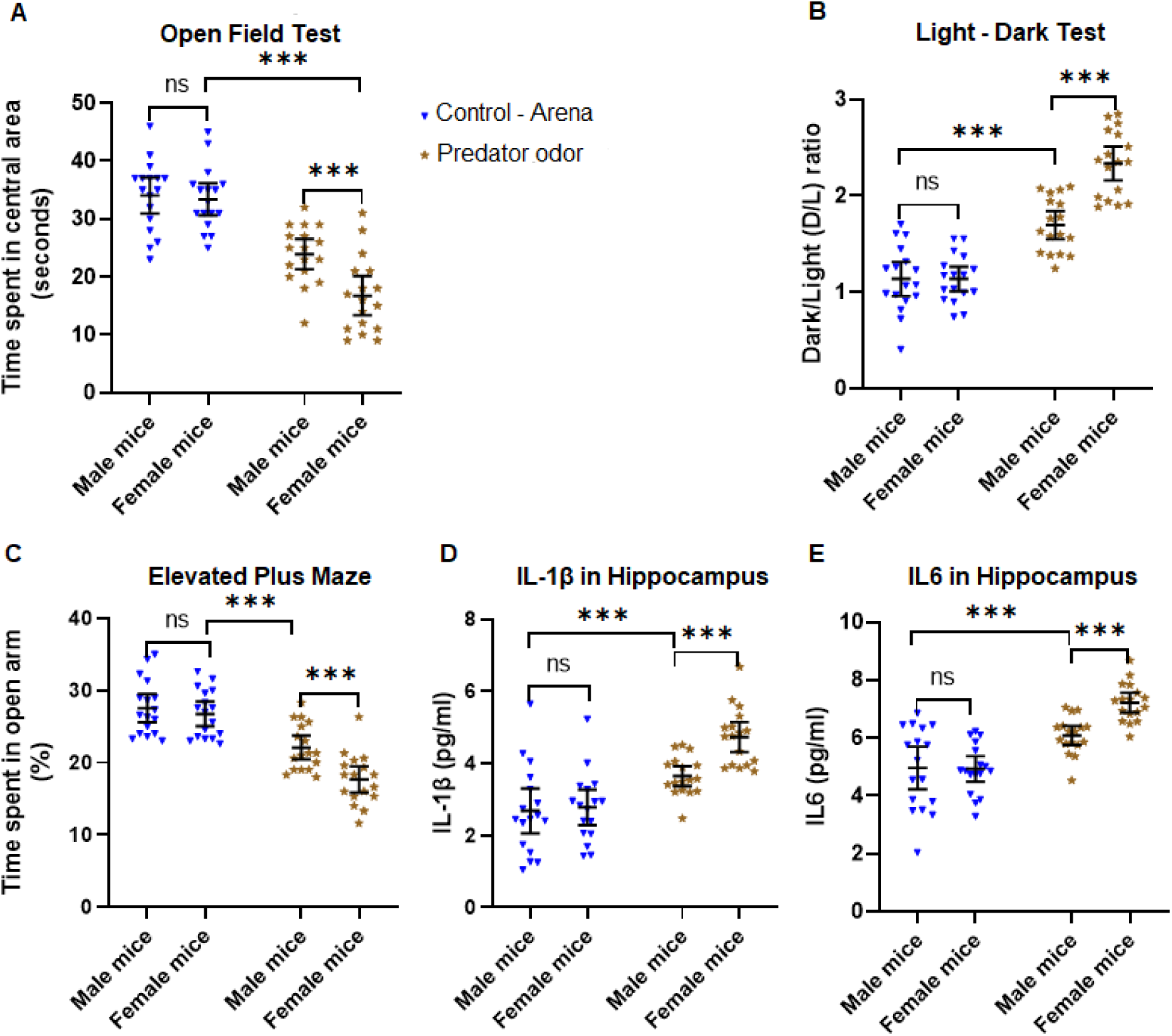
Female mice exhibit exacerbated anxiety following predator odor stress. **(A)** Evaluation of anxious behavior of stressed (predator odor) and non-stressed control mice using open field test (OFT). Female mice exposed to predator odor exhibited significantly higher anxiety levels when compared to stressed males and controls, as depicted by their hesitation to explore the central area of the OFT. **(B)**The light-dark test showed female mice exposed to predator odor exhibited significantly higher anxiety levels (higher Dark-Light ratios) when compared to the male stressed mice and control, as demonstrated by their reluctance to spend more time in the light chamber of the LDT. **(C)** Elevated plus maze test showed female mice subjected to predator odor displayed significantly increased anxiety levels compared to the other groups, as evidenced by their avoidance to spend more time in the open arms of the EPM. **(D)** Evaluation of IL1β levels by ELISA in hippocampal homogenates revealed female mice exposed to predator odor showed aberrantly higher IL1β as compared to stressed males. **(E)** Exposure of predator odor to female mice caused aberrantly higher IL6 levels in the hippocampus as compared to their male counterparts. All data are presented as Mean with 95% CI (n=17/group); ***p<0.001, ns (not significant); one-way *ANOVA* (Shapiro-Wilk’s, *p*>0.05) followed by Bonferroni *post-hoc* test.

**Figure S2.**
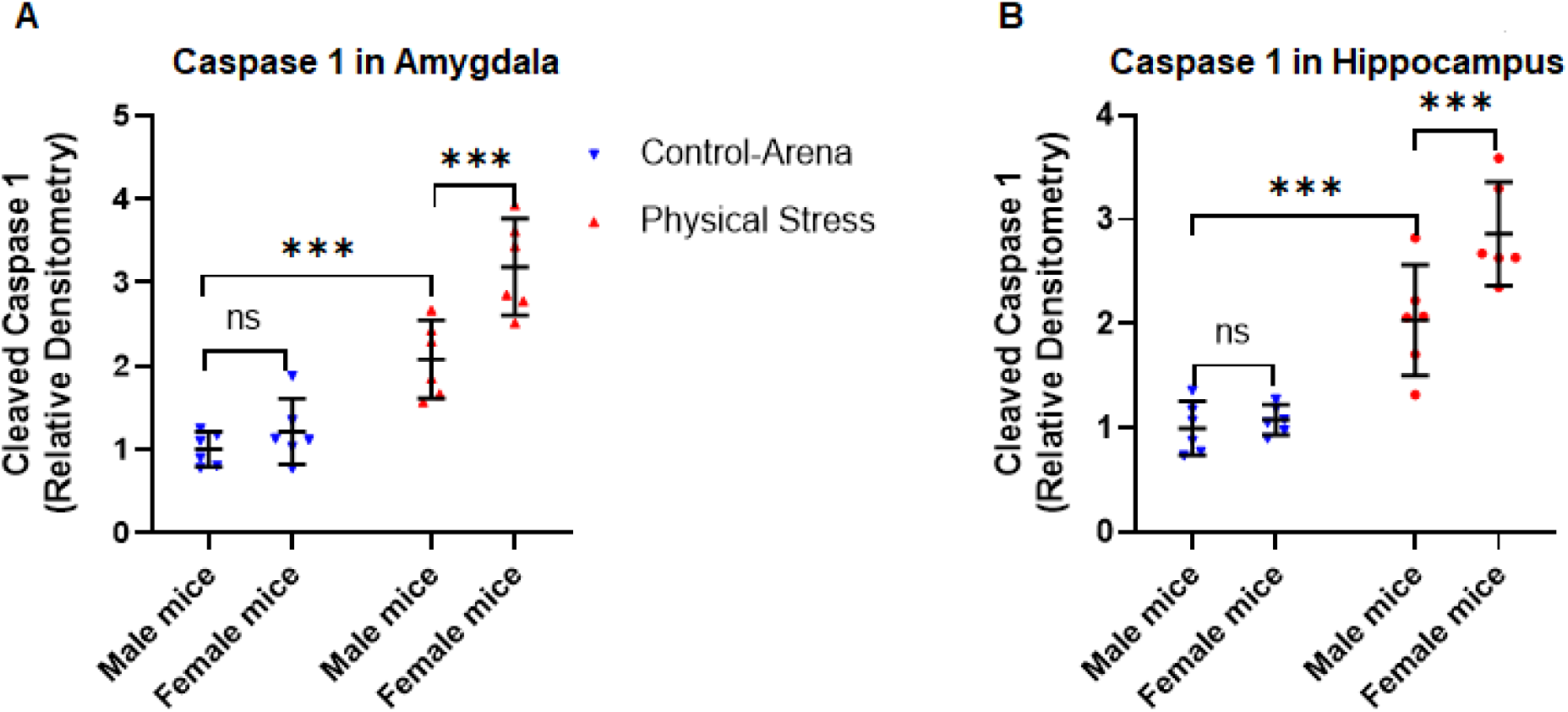
Physical stress in mice leads to induction of cleaved Caspase 1 in amygdala and hippocampus. Physical stress was induced by subjecting mice to restraint and underwater trauma. **(A)** Analysis of amygdala homogenates for cleaved Caspase 1 (p20) using quantitative densitometry of immunoblots. **(B)** Relative densitometry analysis of cleaved Caspase 1 immunoblots from hippocampal homogenates. Mice subjected to restraint and underwater trauma show upregulation of Caspase 1 activity, as evident by the increased level of cleaved Caspase 1 in both amygdala and hippocampus of stressed mice. All data are presented as Mean with 95% CI (n=6/group); ***p<0.001, ns (not significant); one-way *ANOVA* (Shapiro-Wilk’s, *p*>0.05) followed by Bonferroni *post-hoc* test.

**Figure S3.**
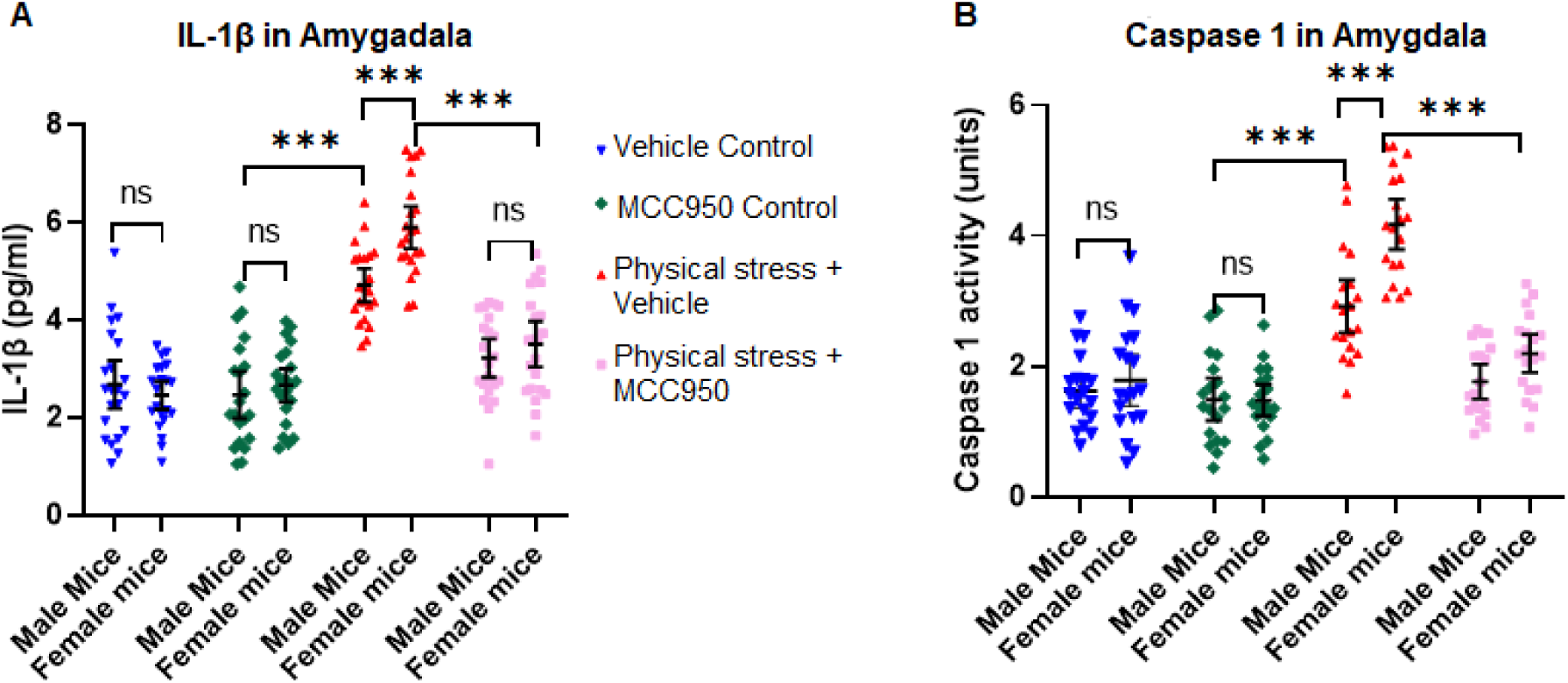
Inhibition of NLRP3 inflammasome by treating stressed mice with MCC950 attenuates IL1β and Caspase 1 activity in amygdala. Physical stress was induced by subjecting mice to restraint and underwater trauma. Control mice were treated with either vehicle or MCC950. Similarly, stressed mice were treated with either vehicle or MCC950. **(A)** Analysis of IL1β in amygdala homogenates by ELISA revealed mice exposed to physical stress showed aberrantly higher IL1β compared to control mice. Treatment of stressed mice with MCC950 showed a significant reduction of IL1β. **(B)** Caspase 1 activity in amygdala: intraperitoneal administration of MCC950 in physically stressed mice showed reduced Caspase 1 activation, elucidating key role of NLRP3 in anxiogenic Capsase 1 - IL1β pathway. Caspase 1 activity in amygdala homogenates was measured by Caspase 1 activity assay kit. All values are presented as Mean with 95% CI (n=19-22/group); ***p<0.001, ns (not significant); two-way ANOVA (Shapiro-Wilk’s, *p*>0.05) followed by Bonferroni *post-hoc* test.

**Figure S4.**
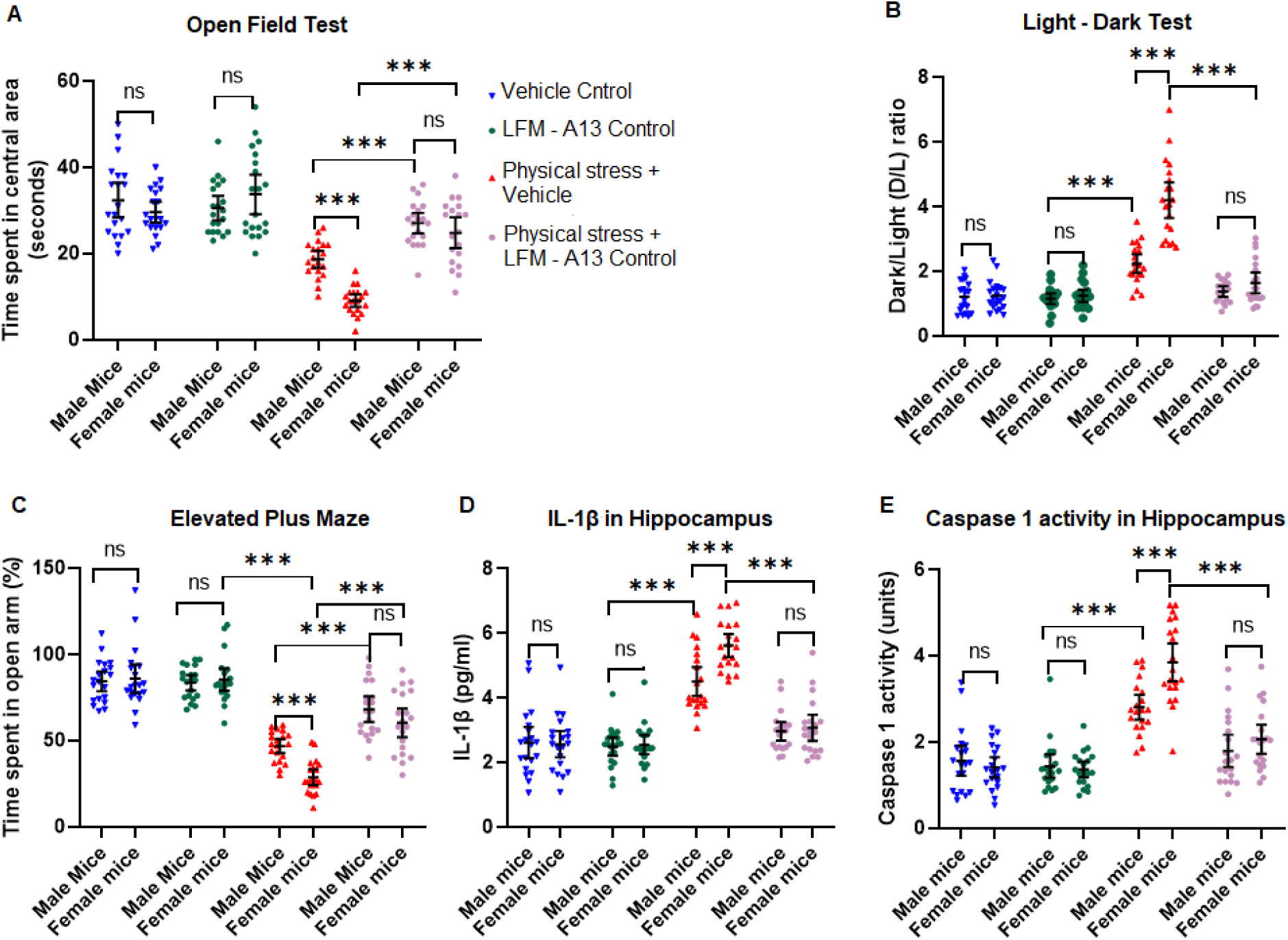
Inhibition of BTK with LFM-A13 in physically stressed mice provided protection from hyper-anxious behavior and inhibited the proinflammatory pathway. **(A)** Analysis of the open field test, following the administration of LMF-A13 in physically stressed mice, there was evidence of significantly decreased anxiety level, as illustrated by the increase in the time of exploration in the central area of the OFT. **(B)** Light-Dark test: physically stressed mice dosed with LMF-A13 showed significant rescue from anxiety, as illustrated by the significant increase in time to spend in the light chamber of the LDT. **(C)** Results of an elevated plus maze also revealed physically stressed mice dosed with LMF-A13 displayed a significant reduction in anxiety, as evidenced by the increase in the duration of time spent in the open arms of the EPM. **(D)** Examination of IL1β levels by ELISA in the hippocampus revealed physically stressed mice dosed with LMF-A13 showed diminished IL1β levels. **(E)**Caspase 1 activity in the hippocampus: physically stressed mice administered with LMF-A13 showed reduced Caspase 1 activation, elucidating significant rescue from anxiogenic proinflammatory pathway. All values are presented as Mean with 95% CI (n=20/group); ***p<0.001, ns (not significant); two-way ANOVA (Shapiro-Wilk’s, *p*>0.05) followed by Bonferroni *post-hoc* test.

